# Single-molecule tracking reveals two low-mobility states for chromatin and transcriptional regulators within the nucleus

**DOI:** 10.1101/2022.07.25.501476

**Authors:** Kaustubh Wagh, Diana A Stavreva, Rikke AM Jensen, Ville Paakinaho, Gregory Fettweis, R Louis Schiltz, Daniel Wüstner, Susanne Mandrup, Diego M Presman, Arpita Upadhyaya, Gordon L Hager

## Abstract

How transcription factors (TFs) navigate the complex nuclear environment to assemble the transcriptional machinery at specific genomic loci remains elusive. Using single-molecule tracking, coupled with machine learning, we examined the mobility of multiple transcriptional regulators. We show that H2B and ten different transcriptional regulators display two distinct low-mobility states. Our results indicate that both states represent dynamic interactions with chromatin. Ligand activation results in a dramatic increase in the proportion of steroid receptors in the lowest mobility state. Mutational analysis revealed that only chromatin interactions in the lowest mobility state require an intact DNA-binding domain as well as oligomerization domains. Importantly, these states are not spatially separated as previously believed but in fact, individual H2B and TF molecules can dynamically switch between them. Together, our results identify two unique and distinct low-mobility states of transcriptional regulators that appear to represent common pathways for transcription activation in mammalian cells.

## INTRODUCTION

The eukaryotic genome is highly organized across several length scales. DNA wraps around nucleosomes to form chromatin, which loops, forms topologically-associating domains (TADs), chromosomes, and chromosome territories (*1*). This organization is crucial for the regulation of gene expression. Genes that are present in more accessible regions or within three-dimensional proximity of their cis-acting elements (enhancers, promoters) are more likely to be expressed (*2*). Transcription factors (TFs) bind consensus motifs within enhancers and promoter-proximal regions, and this binding triggers the recruitment of cofactors, remodelers, co-repressors, or co-activators, all of which act in concert to regulate target genes (*3*). This is a highly dynamic process with TFs only transiently interacting with chromatin on a timescale of seconds (*4-6*). Chromatin itself is a highly dynamic polymer, subject to thermal fluctuations and active forces such as transcription (*7*), loop extrusion (*8*), DNA damage repair, and replication (*9*). How TFs navigate this complex nuclear microenvironment to find their binding sites remains poorly understood.

Over the past decade, single-molecule tracking (SMT) has emerged as a powerful tool to interrogate the dynamics of proteins in living cells. In bacteria, TFs have been shown to undergo a combination of 3D diffusion and 1D facilitated diffusion (sliding) to find their target sites (*10*). Mammalian nuclei present a much bigger challenge to the TF search for relevant motifs since the nucleus contains several levels of organization. Chromatin in mammalian cells exhibits complex dynamic signatures, showing micron-scale coherence on a timescale of 10 s (*11*) and recent SMT studies have found that transcription (*12*) and loop extrusion (*13*) constrain chromatin mobility. Classification of fast TF and histone H2B trajectories into five mobility groups revealed a spatial patterning of mobility states (*14*), with lower mobility states occupying the nuclear periphery and perinucleolar regions, which are typically associated with heterochromatin. Similarly, fast SMT showed that nucleosomes exhibit two mobility states on a timescale of 500 ms, which were then modeled as spatially separate domains of ‘fast’ and ‘slow’ chromatin (*15*). In both these studies, single-molecule trajectories were sampled rapidly (100 Hz (*14*) or 20 Hz (*15*)) and for short times (≤ 500 ms). However, TF dwell times have been shown to obey a power-law distribution, with some binding events lasting for tens of seconds (*16*). To identify mobility states that are important on these timescales, it is essential to study the molecules that remain bound for similar times. Furthermore, chromatin is a viscoelastic polymer showing different dynamic signatures at short and long timescales (*11, 17*). This makes it important to complement these fast SMT studies with SMT studies sampling longer TF binding events to get a more complete picture of chromatin and TF dynamics.

Despite extensive studies of chromatin dynamics, several questions remain open: Which modes of chromatin mobility can we detect at timescales meaningful for TF binding? Are these mobility states spatially separate or can individual nucleosomes switch between them? Do TFs and coregulators exhibit similar mobility states as chromatin? For inducible TFs, how do these states change upon ligand-activation? Which domains of TFs are key determinants of mobility and chromatin interactions?

Here, we use SMT along with a systems-level machine-learning algorithm to address these questions. First, we focus on H2B as a marker for chromatin, and find that H2B exhibits two distinct low-mobility states. Individual H2B molecules dynamically switch between these states, challenging the view that chromatin forms long-lasting and spatially separated mobility domains. Next, we used our analysis framework to study steroid receptors (SRs), which are hormone-inducible TFs. We find that SRs, along with other coregulators, show the same two low-mobility states as H2B, indicating that TF motion is correlated with that of chromatin. Like H2B, TFs and coregulators can also switch between these two states. Upon activation of SRs, the bound fraction as well as the proportion of molecules in the lowest mobility state increase significantly, indicating that this state is associated with the active form of SRs. Focusing on the peroxisome proliferator-activated receptor gamma 2 (PPARγ2), we show that engagement with chromatin in the lowest mobility state requires an intact DNA-binding domain (DBD) as well as domains important for the formation of heterodimeric protein complexes that enhance chromatin binding and transcriptional output. Finally, we discuss our results in the context of recent studies and propose a new model for transcription factor dynamics in mammalian cells.

## RESULTS

### Chromatin mobility is characterized by dynamic switching between two distinct low-mobility states

We performed SMT of cells expressing HaloTag-protein chimeras (with H2B serving as a probe for chromatin) to determine the spatial mobility of proteins. We labeled the HaloTag-protein chimeras with low concentrations (5 nM) of JF_549_ dye (*18*), and imaged cell nuclei using highly inclined laminated optical sheet (HILO) microscopy (*19*) (Fig. 1A). We are most interested in the spatial mobility of bound events that last on the order of tens of seconds as they were shown to be correlated with transcriptional outcomes (*20*). Since photobleaching prevents rapid imaging for long times (*21*), we imaged the cells every 200 ms, with short exposure times of 10 ms to minimize motion blur (Movie S1). Particles were tracked using a custom algorithm (see Methods). A representative temporal projection of an H2B SMT image stack along with particle tracks is shown in Fig. 1B.

**Fig. 1:**
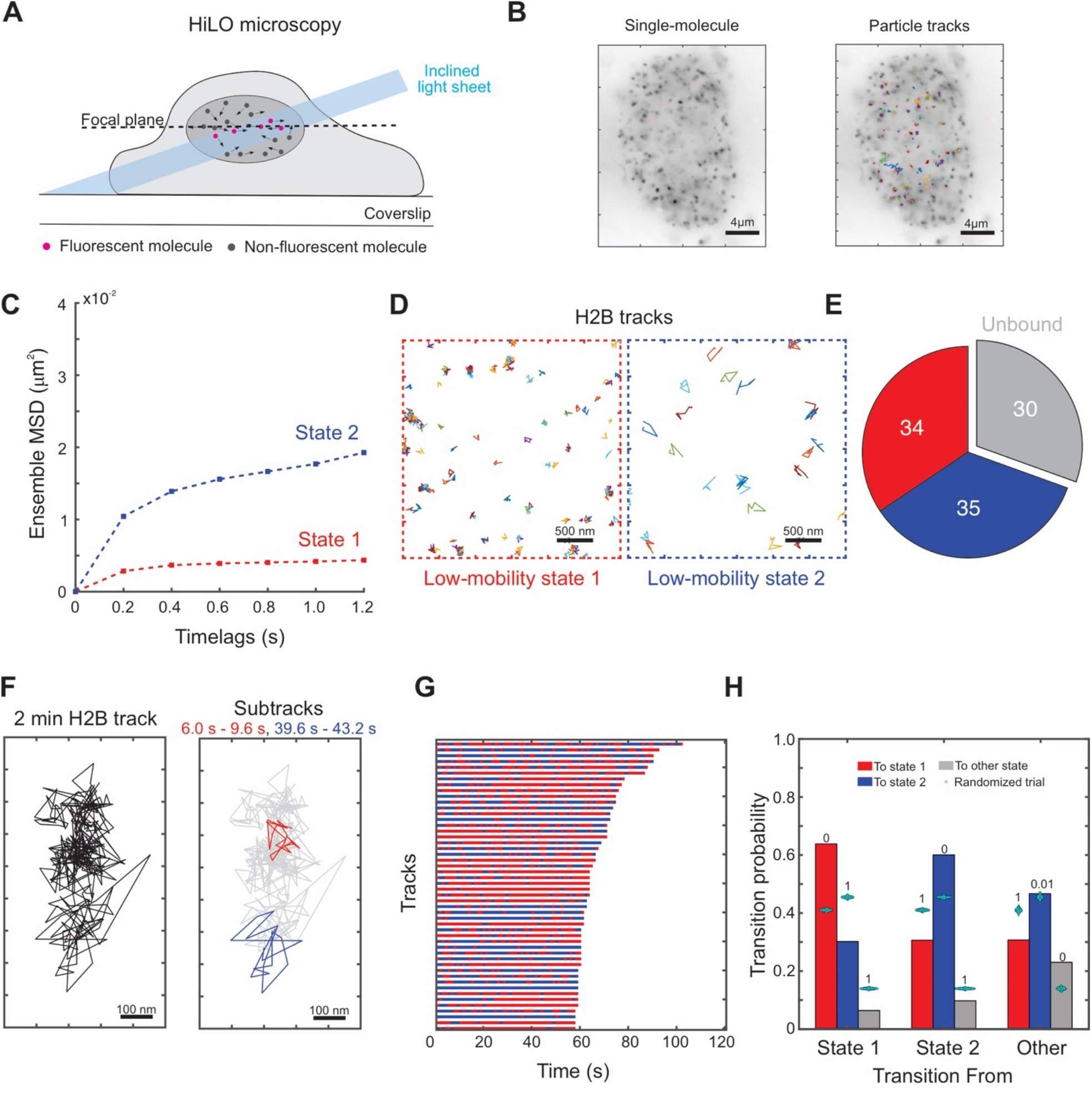
Histone H2B shows two distinct low-mobility states. **(A)** Schematic of single-molecule tracking experiment. **(B)** (left) Time projection of a representative H2B-Halo single-molecule tracking (SMT) movie (right) Overlaid with tracks. **(C)** Ensemble mean-squared displacement (MSD) for histone H2B (N_cells_ = 149, N_tracks_ = 25,298, N_sub-tracks_ = 88,934). Error bars denote the standard error of the mean. **(D)** Sample tracks assigned to low-mobility state 1 (red), low-mobility state 2 (blue) for H2B. **(E)** Piecharts of proportions of H2B sub-tracks assigned to different mobility states. **(F)** (left) Sample H2B track (right) Sub-tracks of length 3.6s color coded by state assignment (state 1 in red and state 2 in blue). **(G)** Temporal reconstruction of the 50 longest tracks for histone H2B. **(H)** Transition probabilities for H2B among states 1, 2, and all other states. Cyan swarmcharts represent transition probabilities calculated from 1000 randomized ensembles. Number above the bars represent the proportion of randomized trials that have a higher transition probability than the respective calculated transition probability.

To quantify and characterize the mobility of H2B, we used a systems-level classification algorithm (perturbation expectation maximization version 2 (pEMv2)) to classify H2B trajectories into different diffusive states (*22*). Given a collection of trajectories without any *a priori* knowledge of the underlying diffusive states, pEMv2 utilizes machine learning along with a Bayesian information (BIC) criterion to uncover a set of diffusive states from a complex distribution of diffusivities. To minimize errors due to transitions within a track, while retaining sufficient numbers of data points for classification, we split our tracks into sub-tracks of length 7-frames (Fig. S1A) (*23*). Since pEMv2 is a probabilistic algorithm, we assign each sub-track to the state for which it has the highest posterior probability, filtering out sub-tracks with similar probabilities of belonging to multiple states (Fig. S1, B and C) (see Methods for details). After classification by pEMv2, we removed any states with a population fraction smaller than 5% (see Methods, Fig. S1, A and C). We then used the ensemble mean-squared displacement (MSD) of these states to compare diffusive states across proteins and conditions. The MSD curve serves as a good metric for the exploration size and diffusivity of an ensemble of particles (*24*).

In adenocarcinoma 3617 cells (*25*), we found that the ensemble of H2B trajectories converged to seven mobility states, but the bulk of sub-tracks were classified into two states (Fig. S1C) based on the posterior probability of assignment to particular states (see Methods). Inspection of the ensemble MSD for both these states (Fig. 1C; henceforth referred to as states 1 and 2), as well as randomly sampled sub-tracks (Fig. 1D) shows that state 2 has a higher exploration radius than that of state 1 and that these states are distinct. States 1 and 2 each account for ∼35% of all sub-tracks while ∼30% of H2B molecules are unbound (Fig. 1E) (see Methods). Our data agree with recent studies (*15, 23*), which showed that H2B exhibits two distinct mobility states. Ashwin et al. attributed these states to distinct *spatially separated* domains of fast and slow chromatin (*15*). However, the bulk of the data in that study represent relatively short tracks that last less than 500 ms (*15*). While each of our sub-tracks is of a comparable length (1.2 s), the parent tracks are longer, with some lasting up to 2 minutes (Fig. 1, F and G).

To determine whether the two mobility states correspond to spatially separated chromatin domains that persist over seconds, we analyzed the dynamics of the two low mobility states within individual tracks. We generated a temporal reconstruction of state dynamics by coloring in sub-tracks by the color of the state they are assigned to (Fig. S1, D and E). If indeed, the two mobility states are spatially separated, we would expect to see entire tracks that belong to state 1 or 2. Strikingly, we found that the same H2B molecule dynamically switches between both low mobility states as shown for a representative track in Fig. 1F. Note that while we have picked spatially separated sub-tracks for ease of visualization (Fig. 1F), state 1 and state 2 sub-tracks overlap throughout the parent track (for example, Fig. S1D). More generally, across an ensemble of the 50 longest tracks, we observed similar switching behavior between these two states (Fig. 1G).

We then quantified the transition probabilities for all tracks that contain at least three sub-tracks (see Methods). To determine whether these transition probabilities are statistically significant, we performed a permutation test: we generated 1000 ensembles of randomly permuted sub-tracks and calculated the transition probabilities for these ensembles (see Methods). This approach has been used previously to test the statistical significance of transition matrices in atmospheric Markov chains (*26*). Our analysis shows that H2B molecules in states 1 and 2 prefer to remain in the same state (Fig. 1H), and that this occurs with a higher probability than would be expected from random ensembles with the same population fractions (Fig. 1H). On the other hand, while we do observe transitions from state 1 to state 2 and vice versa, our permutation test shows that these transitions occur less frequently than in a random ensemble (Fig. 1H).

Together, our data suggest that rather than forming spatially separated domains of higher or lower mobility, chromatin can switch dynamically between these two mobility states. However, this only becomes apparent when we track nucleosomes over longer timescales.

### SRs also exhibit two low-mobility states, with ligand-dependent population fractions

Having established that chromatin has two dynamic mobility states, we turned our attention to TFs. How is TF mobility different from that of H2B? Do active and inactive forms of a TF behave differently? To answer these questions, we applied our analysis framework to study multiple SRs, which are class I nuclear receptors that bind hormone response elements (HREs) as homodimers or homotetramers (*27, 28*). Some SRs, like the glucocorticoid receptor (GR) and the androgen receptor (AR), are predominantly cytoplasmic in the absence of hormone with a small nuclear fraction, while the estrogen receptor (ER) is mostly nuclear (*29*). In the case of the progesterone receptor (PR), it can be either predominantly cytoplasmic or nuclear, depending on isoform (*30*). Agonist binding triggers a conformational change, nuclear translocation (for GR, AR, and PR), oligomerization, and binding to HREs. We tracked unliganded ER, and the small nuclear fraction of unliganded GR, PR, and AR and contrasted these with their corresponding ligand-activated receptors.

All tested SRs, with and without activation by hormone, exhibit two distinct low-mobility states as well as a small population of one or two higher mobility states (Fig. 2, A to I, Fig. S2). Since a majority of the sub-tracks belong to the two low-mobility states (Fig. S2), we will focus on these for the rest of the study. As can be seen qualitatively from sample tracks belonging to these states (Fig. 2A), and quantitatively from ensemble MSD plots (Fig. 2, B to I), these states have different mobility signatures. On comparing these with the states recovered for H2B, we find that all the examined SRs exhibit the same two low-mobility states as H2B (Fig. 2, B to I). This implies that SR mobility states and chromatin dynamics are correlated at our observed timescales.

**Fig. 2:**
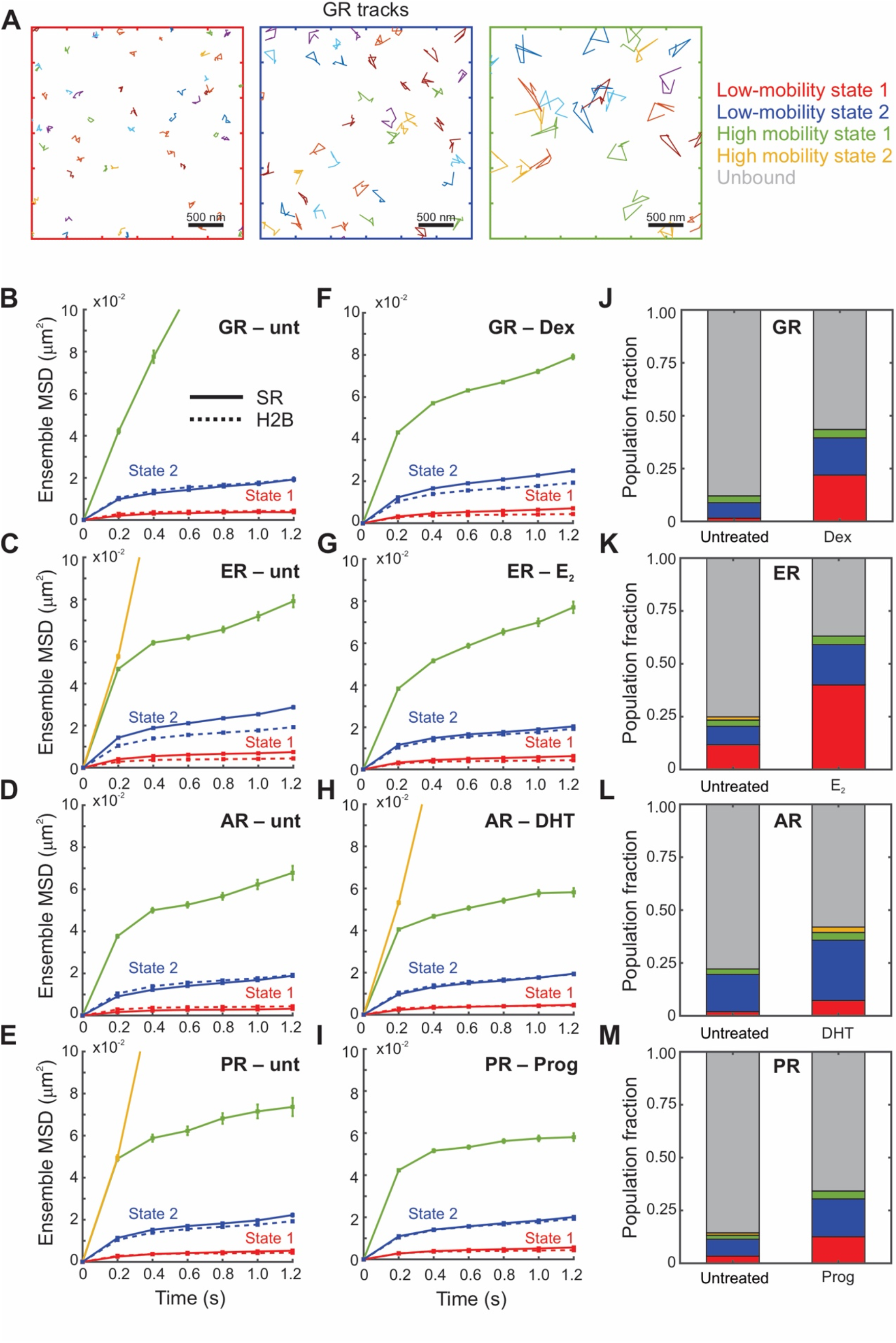
Steroid receptors also exhibit two low-mobility states, with ligand-dependent population fractions. **(A)** Sample tracks for the glucocorticoid receptor (GR). (left) Low-mobility state 1, (middle) low-mobility state 2, (right) High-mobility state. **(B – I)** Ensemble mean-squared displacement for indicated steroid receptor (solid lines) and histone H2B (dashed lines), error bars denote the standard error of the mean: **(B)** Untreated GR (N_cells_ = 35, N_tracks_ = 386, N_sub-tracks_ = 962). **(C)** Untreated estrogen receptor (ER) (N_cells_ = 49, N_tracks_ = 4057, N_sub-tracks_ = 9551). **(D)** Untreated androgen receptor (AR) (N_cells_ = 51, N_tracks_ = 1394, N_sub-tracks_ = 4001). **(E)** Untreated progesterone receptor (PR) (N_cells_ = 37, N_tracks_ = 1371, N_sub-tracks_ = 3197). **(F)** GR activated with dexamethasone (N_cells_ = 238, N_tracks_ = 30,652, N_sub-tracks_ = 81,172). **(G)** ER activated with 17β-estradiol (E_2_) (N_cells_ = 50, N_tracks_ = 8147, N_sub-tracks_ = 24,299). **(H)** AR activated with dihydrotestosterone (DHT) (N_cells_ = 38, N_tracks_ = 5160, N_sub-tracks_ = 12,697). **(I)** PR activated with progesterone (Prog) (N_cells_ = 41, N_tracks_ = 4951, N_sub-tracks_ = 14,899). **(J – M)** Comparative barcharts showing population fractions of various states for the indicated steroid receptors.

To better understand the biological origin of these mobility states for SRs, we compared the population fractions of the states before and after hormone activation. All four SRs show an increase in the overall bound fraction upon activation (Fig. 2, J to M). All SRs show a dramatic increase in the proportion of the lowest mobility state, state 1: 10-fold for GR (Fig. 2J); 3.3-fold for ER (Fig. 2K); 3.5-fold for AR (Fig. 2L); and 4-fold for PR (Fig. 2M). These are accompanied with a smaller increase in the relative proportion of state 2: 2.5-fold (GR), 2.1-fold (ER), 1.5-fold (AR), and 2.2-fold (PR) (Fig. 2, J to M). It should be noted that 3617 cells do not express endogenous AR and PR (*31, 32*), and therefore may not provide a native chromatin context for AR and PR binding. This likely results in the relatively low population fractions observed for state 1 in these cells (Fig. 2, L and M). Taken together, these data suggest that state 1, the lowest mobility state of SRs, better correlates with their activation status than state 2, which implies that either binding of activated SR to chromatin constrains its mobility and/or activated SRs are more capable of interacting with chromatin in state 1.

pEMv2 is a systems-level analysis that produces discrete mobility states and posterior probability distributions that maximize a defined log-likelihood function (*22*). We used an alternative method to test the generality of our observed mobility states. Given a collection of trajectories, we can calculate the van Hove correlation (vHc) function or step-size distribution. The calculated vHc can then be approximated as a superposition of Gaussian basis functions (see Methods), from which we can iteratively calculate the distribution of MSDs that gives rise to the calculated vHc. We used the iterative algorithm developed by Richardson (*33*) and Lucy (*34*) and successfully implemented it to study nucleosome dynamics (*15*) and to calculate the distribution of MSD (or equivalently the diffusivity distribution). We refer to this analysis as ‘RL analysis’ in the rest of the manuscript. We find once again that the predicted MSD distribution for H2B, shown here for a time lag of 0.8 s (Fig. S3, A and B) has two main populations, confirming the two states recovered from pEMv2. The bimodal distribution of MSDs was observed for other time lags (0.6 s - 1.2 s) as well (not shown), indicating the generality of our findings. Similar analysis for hormone-activated SRs (GR, ER, AR, PR) also showed two distinct low-mobility states supporting our pEMv2 results (Fig. S3, C to J). Consistent with the thinly populated higher mobility states detected by pEM, we observe some higher mobility states in the distribution of MSDs as well (Fig. S3).

### SRs dynamically switch between the two low-mobility states

Since we observe H2B molecules switching between the two low-mobility states, we next examined whether SRs also exhibited similar switching behaviors. Visual inspection of tracks showed that the same Dex-activated GR molecule could switch between these two mobility states (Fig. 3, A and B), with the state 2 sub-tracks exhibiting larger jumps (Fig. 3B). We then compared the switching behavior of SRs before and after hormone stimulation.

**Fig. 3:**
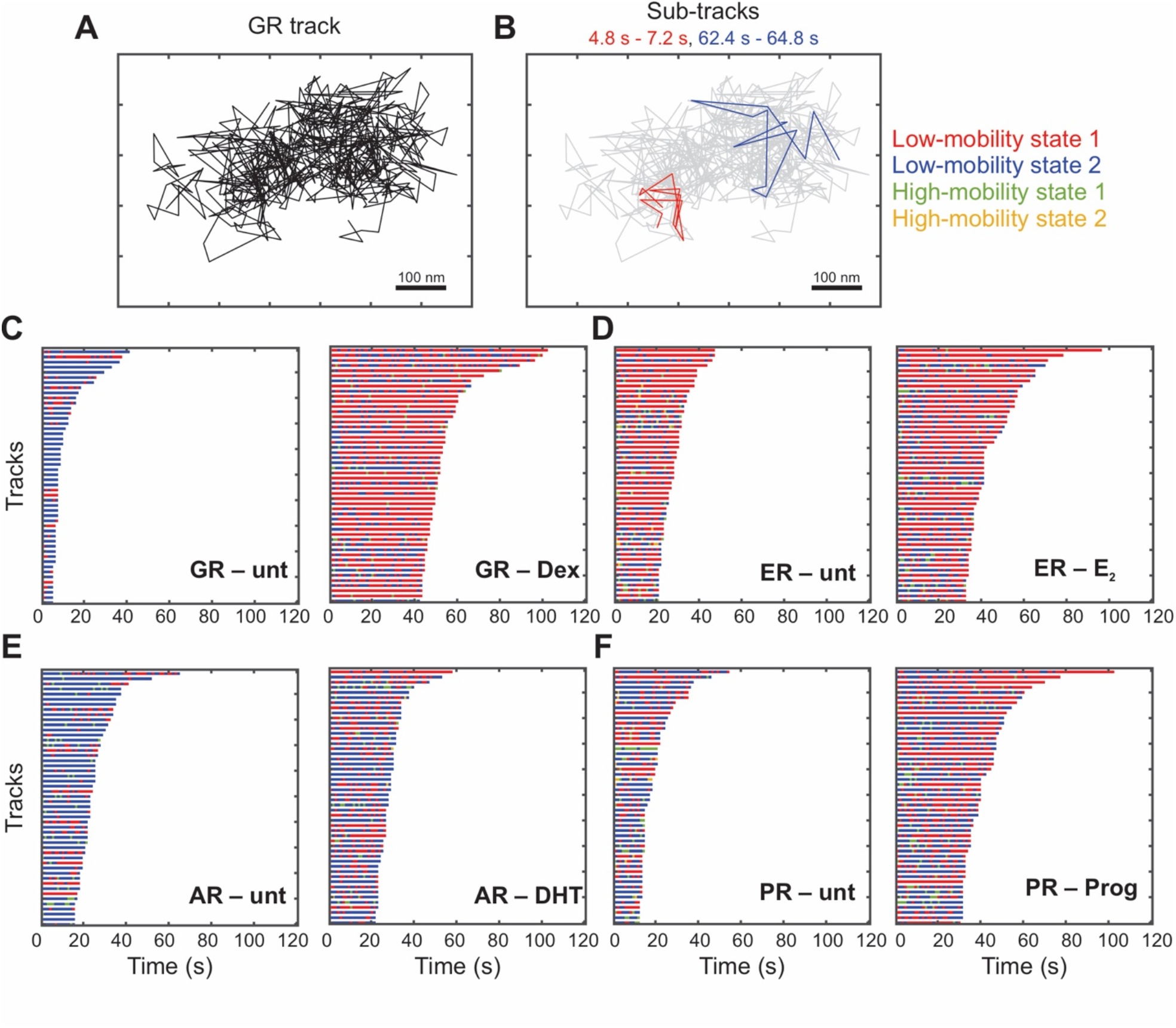
Transcription factors dynamically switch between two low-mobility states. **(A)** Sample GR track. **(B)** Sub-tracks of length 2.4s from the same track as in **(A)** color-coded by state assignment (state 1 in red and state 2 in blue). **(C – F)** Temporal reconstruction for the 50 longest tracks for steroid receptors. (left) without hormone (right) upon activation by hormone. **(C)** Glucocorticoid receptor. **(D)** Estrogen receptor. **(E)** Androgen receptor. **(F)** Progesterone receptor.

Upon activation by hormone, we observed an increase in overall dwell times (indicated by larger track durations of the longest tracks) of all SRs, as has been reported previously (*21*) (Fig. 3, C to F). We find that similar to H2B, SRs also exhibit switching between the two lowest mobility states (Fig. 3, C to F). In addition to showing the same ensemble MSD (Fig. 2, B to I), TFs and H2B both exhibit dynamic transitions between the two low-mobility states (Fig 1, F to H, and Fig. 3, C to F), supporting the hypothesis that the low-mobility states represent different modes of chromatin engagement. Quantifying the probability of transitions between these states, we observed that GR (Fig. S4, A and B), ER (Fig. S4, C and D), and PR (Fig. S4, E and F) molecules in states 1 and 2 prefer to remain in the same state, while transitions into state 2 are dominant for AR (Fig. S4, G and H). Ligand activation results in an ∼13% increase in state 1 to state 1 transitions for GR and a corresponding 6% increase for ER (Fig. S4, A to D), while AR and PR show very subtle differences with and without agonist (Fig. S4, E to H). For unliganded SRs, state 2 to state 1 transitions are not significantly different at the 99% confidence level from those obtained for ensembles of random permutations (Fig. S4, A, C, E, G) (see Methods). However, upon ligand activation, these transitions occur with a higher probability than corresponding transitions for unliganded SRs (Fig. S4) but occur less frequently than the random ensemble. These data suggest that activation of SRs by hormone results in an increase in transitions into state 1 and that these transition probabilities are significantly different from those in a random ensemble.

Collectively, our data show that SRs exhibit two distinct mobility states, which correlate with the mobility states of chromatin. Further, SRs and histones frequently switch between these states, underscoring the fact that these states are not spatially separated. Ligand activation dramatically increases the population fraction of state 1. Taken together, these imply that state 1 represents a mobility state that is important for hormone-mediated gene regulation.

### Other transcriptional regulators also exhibit two distinct mobility states

Since we observed two distinct mobility states for SRs, which represent different modes of chromatin engagement, we hypothesized that other transcriptional regulators should also exhibit these two states. To test this hypothesis, we performed SMT experiments and subsequent analysis on several nuclear proteins with different functions.

RELA/p65 is an important subunit of the NF-κB transcription factor, which is activated in response to many external stimuli (*35*). Glucocorticoid receptor-interacting protein 1 (GRIP1), also known as nuclear receptor coactivator 2 (NCoA2) is a coregulatory protein that is recruited to DNA by nuclear receptors in response to ligand-activation (*36*). GRIP1 facilitates nuclear receptor-mediated gene regulation by acetylating histone tails, thereby modulating chromatin accessibility (*36*). Mediator of RNA polymerase II transcription subunit 26 (MED26) is a subunit of the Mediator complex that assists RNA polymerase II-mediated transcription by recruiting accessory proteins that promote transcriptional elongation (*37*). SMARCA4 (also known as BRG1) is an ATP-dependent remodeler that is a part of the SWI/SNF complex. SMARCA4 modulates gene expression by changing chromatin accessibility through its remodeling activity (*38*). CCCTC-binding factor (CTCF) is important for 3D genome organization, leading to the formation of enhancer-promoter loops and regulating the structure of topologically-associating domains (*1*).

For this diverse set of transcriptional proteins with widely varying functions, we observed two qualitatively similar low-mobility states as histone H2B (Fig. 4, Fig. S5). As seen with SRs, these transcriptional regulators also switch between the two low-mobility states (Fig. S6), with molecules preferentially transitioning to the same state (Fig. S6), except RELA and SMARCA4, which show a slight preference to switch from state 1 to state 2 (Fig. S6, A and D, right). These data suggest that all detected TF and coregulator dynamics correlate with the mobility of the local chromatin environment.

**Fig. 4:**
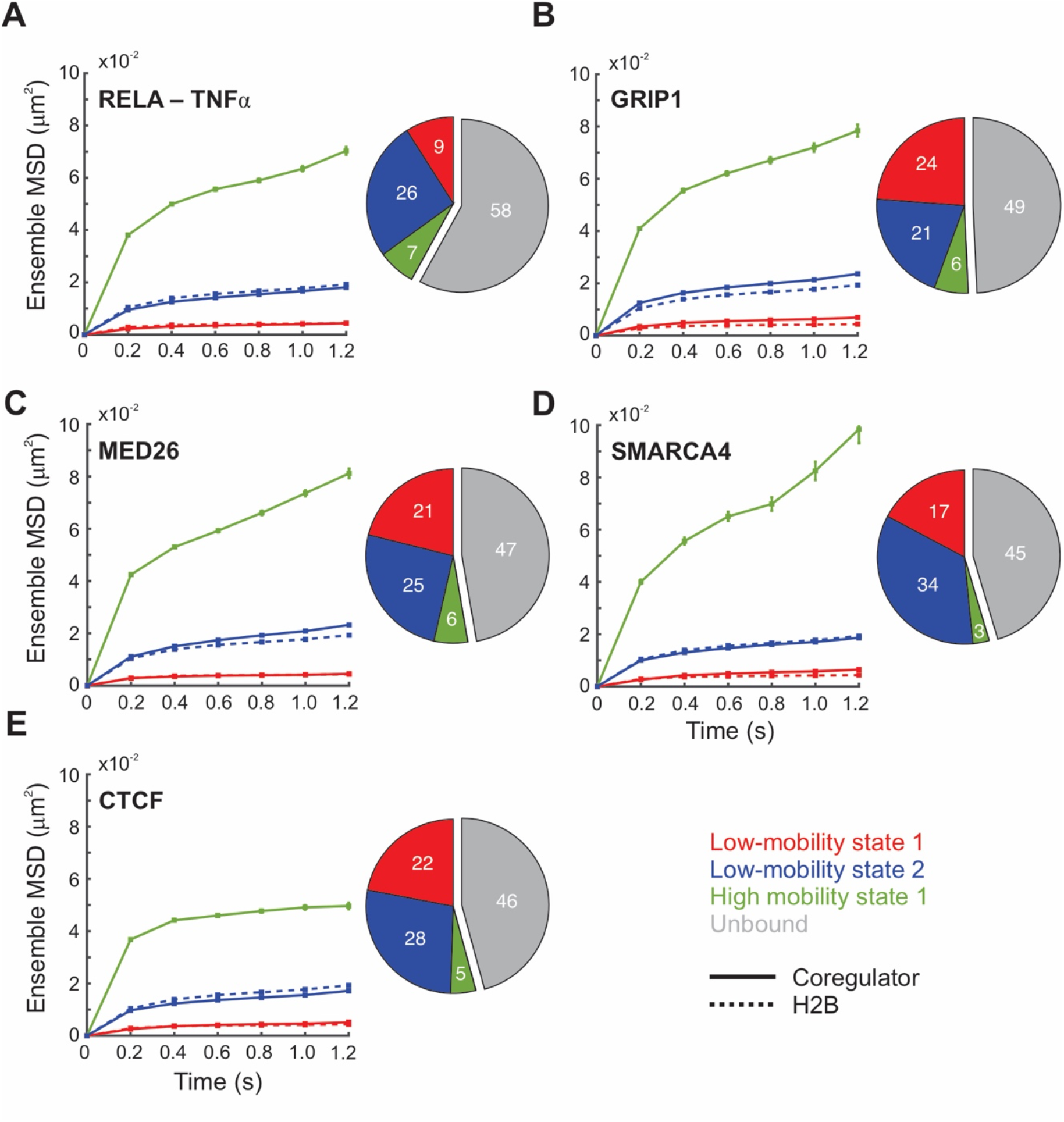
Other transcriptional regulators also exhibit two distinct low-mobility states. **(A – E)** For indicated coregulators (left) Mean-squared displacement plots for indicated transcriptional coregulator (solid lines) and histone H2B (dashed lines), error bars denote the standard error of the mean; (right) piecharts indicating proportions for the various detected mobility states. **(A)** RELA/p65 activated with TNFα (N_cells_ = 67, N_tracks_ = 9524, N_sub-tracks_ = 24,634). **(B)** GRIP1 (N_cells_ = 36, N_tracks_ = 4847, N_sub-tracks_ = 14,010). **(C)** MED26 (N_cells_ = 57, N_tracks_ = 11,429, N_sub-tracks_ = 29,085). **(D)** BRG1/SMARCA4 (N_cells_ = 22, N_tracks_ = 3179, N_sub-tracks_ = 8112). **(E)** CTCF (N_cells_ = 69, N_tracks_ = 10,457, N_sub-tracks_ = 34,503).

### State 1 of the PPARγ2 requires an intact DNA-binding- and oligomerization-domain

To understand the factors that determine the partitioning of TFs into the two mobility states, we focused on the peroxisome proliferator-activated receptor gamma 2 (PPARγ2), which is a class II nuclear receptor that binds chromatin as a heterodimer with retinoid X receptors (RXR) (*39*) (Fig. 5A, right (inset)). In particular, the existence of well characterized interacting partners and DNA-binding and heterodimerization mutants allows for a systematic study of PPARγ2’s mobility states. We chose 3T3-L1 mouse pre-adipocytes as our model cell line to study PPARγ2 because PPARγ2 is functionally important for adipogenesis (*40, 41*). This allows us to study a TF with functional relevance in its native chromatin context.

**Fig. 5:**
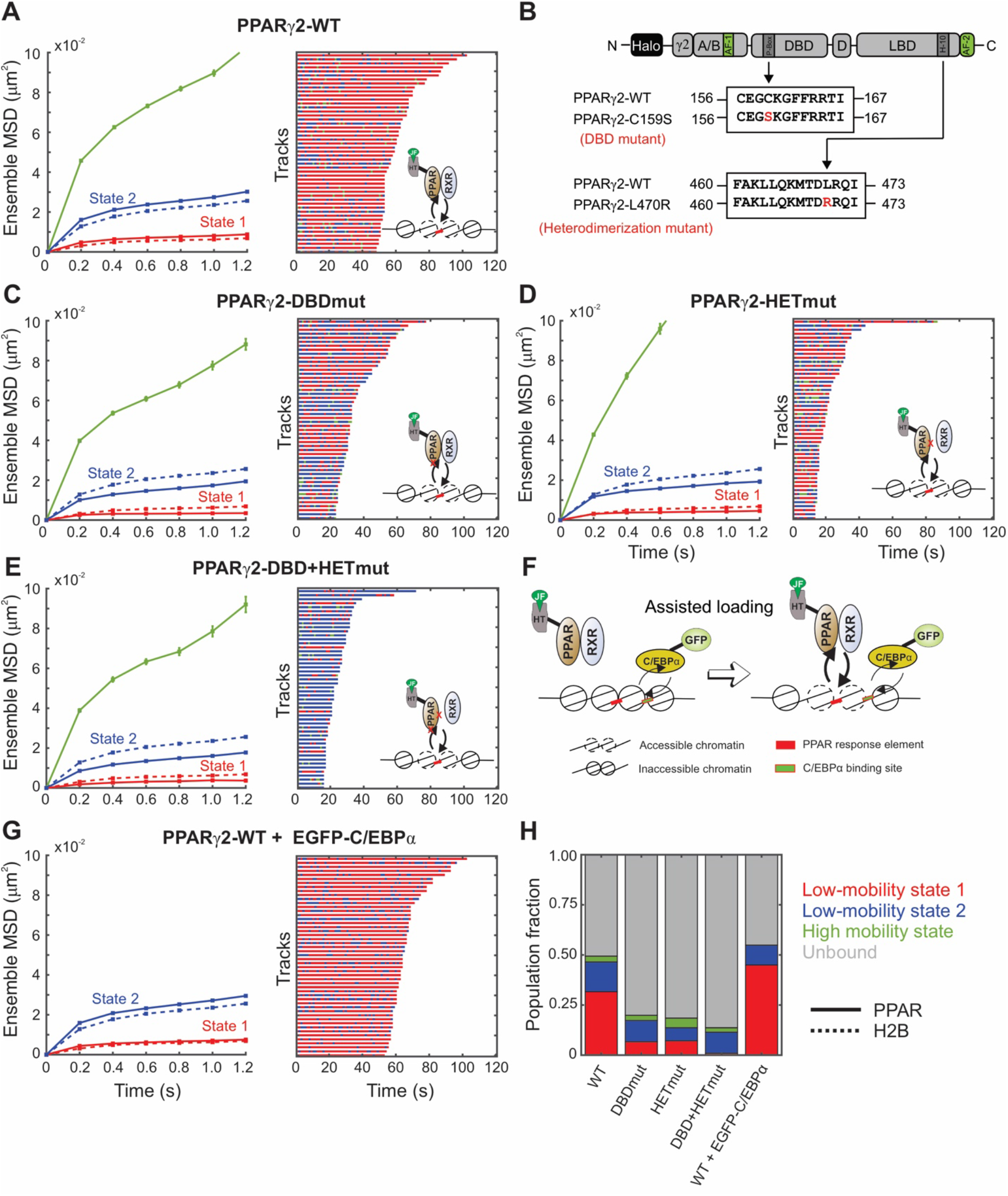
State 1 for PPARγ2 requires intact DNA-binding domain and the ability to form heterodimeric complexes. **(A)** (left) Ensemble mean-squared displacement (MSD) of H2B (dashed lines, N_cell_ = 54, N_tracks_ = 8522, N_sub-tracks_ = 29,262) and wildtype PPARγ2 (solid lines, N_cells_ = 127, N_tracks_ = 20,983, N_sub-tracks_ = 62,848), error bars indicate the standard error the mean (SEM); (right) Temporal reconstruction of the 50 longest tracks along with (inset) a cartoon depicting PPARγ2 binding to PPAR response elements (PPRE). **(B)** Schematic of point mutations to abrogate the DNA-binding domain and heterodimerization domains of PPARγ2. **(C – E)** (left) Ensemble MSD for indicated PPARγ2 mutant. Error bars denote the SEM; (right) Temporal reconstruction of the 50 longest tracks colored by state assignment: **(C)** PPARγ2-DNA-binding domain mutant (PPARγ2-DBDmut) (N_cells_ = 38, N_tracks_ = 3721, N_sub-tracks_ = 9872), **(D)** PPARγ2-heterodimerization mutant (PPARγ2-HETmut) (N_cells_ = 28, N_tracks_ = 1728, N_sub-tracks_ = 4049), **(E)** PPARγ2-DBD + HET mutant (N_cells_ = 46, N_tracks_ = 1695, N_sub-tracks_ = 4046). **(F)** Assisted loading model for C/EBPα mediated PPARγ2 loading. **(G)** (left) Ensemble MSD for PPARγ2-WT with overexpression of GFP-C/EBPα (N_cells_ = 89, N_tracks_ = 18,912, N_sub-tracks_ = 63,842), error bars denote the SEM; (right) Temporal reconstruction of the 50 longest tracks. **(H)** Comparative population fractions for all PPARγ2 variants.

PPARγ2 is one of two PPARγ isoforms expressed from the PPARG gene. PPARγ2 contains 30 additional amino acids on its N-terminal end as compared to PPARγ1 (Fig. 5B). PPARγ1 is expressed in almost all tissues but PPARγ2 is predominantly found in adipose tissue and is important for adipocyte differentiation, fatty acid storage, glucose metabolism, and is a known therapeutic target for diabetes (*42, 43*). During adipogenesis, PPARγ2 and CCAAT enhancer-binding protein alpha (C/EBPα) act in concert to regulate genes essential for the process (*44*).

We first transiently expressed HaloTag-fused H2B and PPARγ2 chimeras in 3T3-L1 cells, performed SMT and analyzed the data with the above-described workflow. As observed in 3617 cells, PPARγ2 and H2B exhibit two distinct and overlapping low-mobility states (Fig. 5A, left). Both PPARγ2 and H2B in 3T3-L1 cells exhibit switching between the two lowest mobility states as seen for other TFs and H2B (Fig. 5A, right, S7, A and B). While H2B molecules in both state 1 and state 2 preferentially transition to the same state (Fig. S7A), PPARγ2 molecules in state 1 remain in state 1 ∼70% of the time but show an equal transition probability from state 2 into both states 1 and 2 (Fig. S7B).

To test the role of the DNA-binding domain (DBD) and the heterodimerization domain (HET) in the two low-mobility states, we first mutated the 159^th^ cysteine to a serine (C159S, henceforth referred to as DBDmut), which has been shown to disrupt the zinc finger and prevent sequence-specific chromatin interactions in vitro (*45*) (Fig. 5, B and C (right, inset)). Disruption of the DBD results in a dramatic reduction in the overall bound fraction and particularly, the population fraction of state 1 as compared to that of wildtype PPARγ2 (Fig. 5, C and H). However, we do not completely lose the bound fraction or the binding in state 1 (Fig. 5, C and H). This is consistent with previous studies that showed that RXR binding to the 3’ half-site of PPAR response elements is more important than PPARγ2 binding to the 5’ half-site for the PPARγ2:RXR complex to stabilize engagement with chromatin (*45*). We also observed an ∼18% increase in the probability of state 2 molecules to remain in state 2, along with a concomitant decrease of ∼19% in the state 2 to state 1 transition probability (Fig. S7, B and C). This suggests that the DBD is important for PPARγ2 to transition from state 2 to state 1.

Mutation of the 470^th^ leucine to arginine (L470R, henceforth referred to as HETmut) eliminates the heterodimerization interface with RXR (*46, 47*) (Fig. 5, B and D (right, inset)). We analyzed this construct to find very similar results as those obtained for the DBD mutant. The overall bound fraction was much smaller than that of PPARγ2-WT, but the same as that of PPARγ2-DBDmut (Fig. 5H). The relative proportion of state 1 was also similar to that of PPARγ2-DBDmut (7%) indicating that monomeric PPARγ2 is still capable of interacting with chromatin, potentially through its intact DBD (Fig. 5H).

By introducing both the DBD and HET mutations simultaneously (PPARγ2-DBD+HETmut; Fig. 5, B and E (right, inset)), we observed that the PPARγ2-DBD+HETmut has an even smaller bound fraction and a vanishingly small proportion of state 1 as compared to those for PPARγ2-WT (Fig. 5, E and H). Like PPARγ2-DBDmut, PPARγ2-HETmut has an impaired ability to transition from state 2 to state 1 (Fig. S7D). As compared to PPARγ2-WT, PPARγ2-DBD+HETmut shows a 31-40% decrease in transitions into state 1 and a 34-38% increase in transitions into state 2 (Fig. S7E). PPARγ2-DBD+HETmut molecules preferentially switch to state 2 from all states (Fig. S7E). Since we have seen that an increase in the proportion of state 1 along with increased transitions into state 1 (from both states 1 and 2) are associated with the active form of SRs (Figs. 2, 3, S4), these data also support the hypothesis that TF engagement with chromatin in state 1 correlates with transcriptional activity. Since these mutations reduce the ability of PPARγ2 to interact with chromatin, we also tested the opposite perturbation: what happens to the two states if we facilitate PPARγ2 binding?

C/EBPα and PPARγ2 have been shown to participate in dynamic assisted loading at closed chromatin sites by recruiting remodelers (*41*) (Fig. 5F). To further test our hypothesis, we over-expressed GFP-fused C/EBPα, which should promote PPARγ2-chromatin interactions in state 1 (Fig. 5F). Consistent with our hypothesis, over-expression of C/EBPα resulted in an increase in the overall bound fraction of PPARγ2 (Fig. 5H) and a 1.4-fold increase in the proportion of state 1 (Fig. 5H). In contrast to the PPARγ2-DBD+HETmut data, over-expression of GFP-C/EBPα results in a 9-16% increase in transitions into state 1 along with an 11-17% decrease in transitions to state 2, with all states showing a preference to switch to state 1 (Fig. S7F). Taken together, our data indicate that binding in state 1 requires an intact DBD and heterodimerization domain, and that this state is correlated with the active form of the TF.

### Tracks with different exploration radii exhibit different switching characteristics

After analyzing sub-tracks using pEMv2, we found that all tested molecules dynamically switch between two low-mobility states. We then used the Richardson-Lucy (RL) analysis to confirm that two states can be recovered from the calculated vHc function (Fig. S3). Since the RL analysis produces a distribution of MSDs, we can use the minima in the MSD distribution to classify entire trajectories into lower or higher mobility populations (Fig. S8). We can then separately analyze the transitions between pEMv2 states for tracks with different overall mobilities (as measured by their MSD at 0.8 s; see Fig. S8).

Analysis of these populations revealed that molecules which were overall less mobile (i.e., with MSD at 0.8 s time lag lower than 0.0075 μm^2^) were predominantly in state 1 (Fig. 6, A to C). Molecules with an overall higher mobility (i.e., with MSD at 0.8 s time lag between 0.0075 and 0.028 μm^2^) exhibited appreciable fractions of state 1 (Fig. 6, D to F). Molecules with overall lower mobility preferentially transition to state 1 (Fig. 6, G to I), while those with an overall higher mobility exhibit a significantly higher probability of switching between these two states (Fig. 6, J to L). This can also be seen by comparing the population fractions of the different mobility states within the two cohorts (Fig. 6M). Combining track-level and sub-track-level analyses thus provide a powerful tool to distinguish between persistent and transient engagement with state 1.

**Fig. 6:**
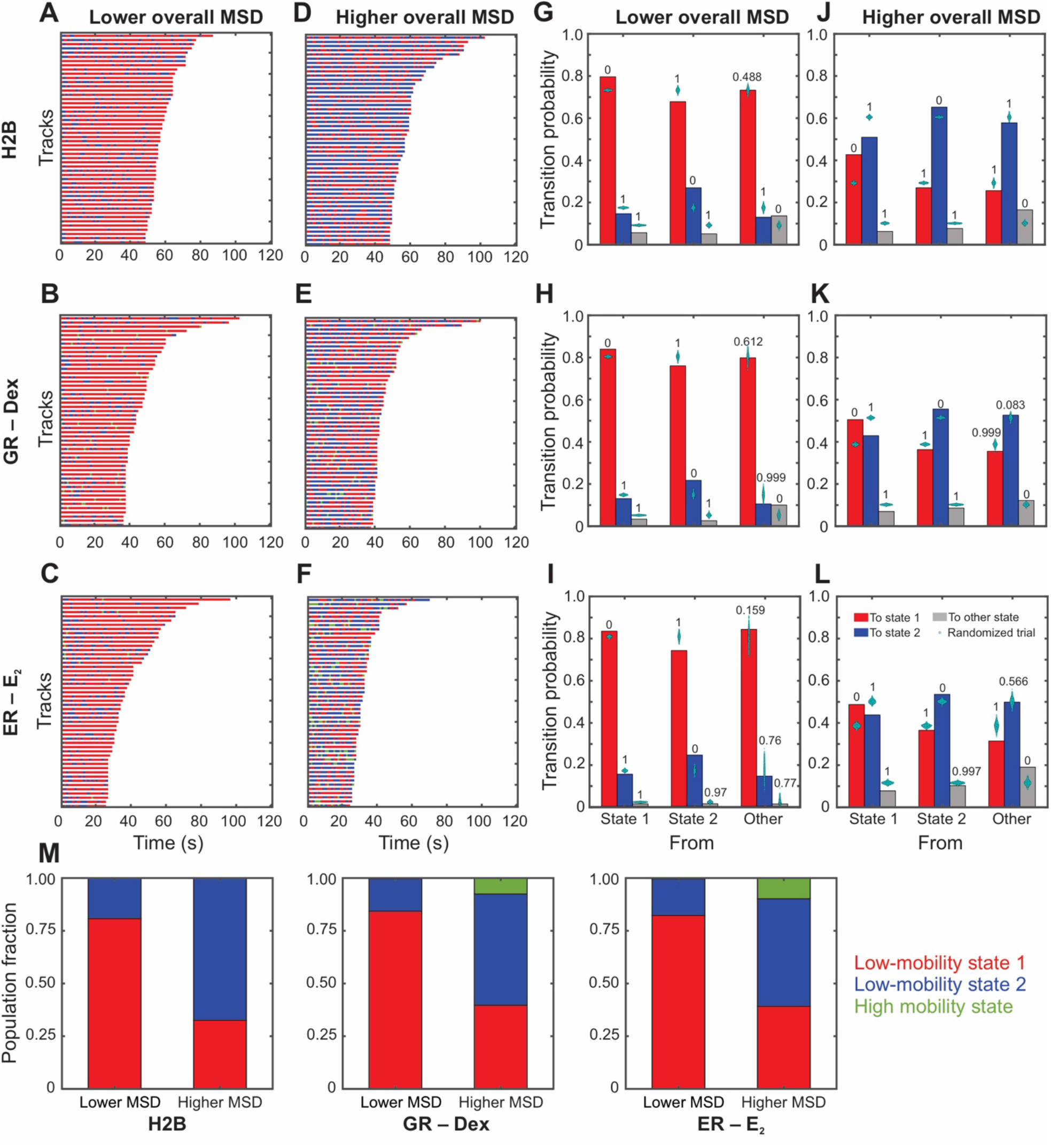
Tracks with different exploration radii exhibit distinct switching patterns. **(A – F)** Temporal reconstruction of the 50 longest tracks of single molecules with **(A – C)** overall lower mobility and **(D – F)** overall higher mobility. The tracks are color-coded to show the pEM identified states of the 1.2 s segments making up entire track. State 1 is depicted in red and state 2 in blue. Higher mobility states are colored green and yellow: **(A, D)** H2B. **(B, E)** Glucocorticoid receptor (GR) activated with Dex. **(C, F)** Estrogen receptor (ER) activated with 17β-estradiol (E_2_). **(G – L)** Transition probabilities calculated for molecules that are less mobile overall **(G – I)** and more mobile **(J – L)**. Transitions into pEM state 1 are shown in red, those into state 2 are shown in blue, and others in gray. Cyan swarmcharts show the results of the transition probability calculation for 1000 randomly permuted ensembles. Numbers above the bars display the proportion of these trials with a transition probability higher than the respective calculated transition probability: **(G, J)** H2B, **(H, K)** GR activated with Dex. **(I, L)** ER activated with E_2_. **(M)** Fraction of segments in pEM state 1 (red), pEM state 2 (blue) and pEM state 3 (green) for trajectories classified in lower and higher mobility states for indicated species.

## DISCUSSION

Single-molecule tracking is a powerful technique to study intranuclear dynamics of individual proteins at the nanoscale with high temporal resolution. Here, using SMT along with a machine learning-based classification algorithm, we identify two distinct low-mobility states for histone H2B (Fig. 1). Previous studies have also found multiple mobility states for H2B (*14, 15*). However, unlike (*14*), our model is not constrained to a fixed number of states. We allow our algorithm to explore up to 15 different states and find that only two states meet our statistical criteria (Fig. S1C). Moreover, unlike the other report which studies dynamics up to 500 ms (*15*), we examine longer timescales on the order of tens of seconds and up to two minutes. Even though we analyze 1.2 s sub-tracks using pEMv2, tracking the same molecule over longer times allows us to identify hitherto hidden transitions between the two low mobility states. We find that unlike previous models (*15*), H2B does not form spatially separated domains of ‘fast’ and ‘slow’ chromatin. Instead, H2B molecules dynamically switch between the two low mobility states (Fig. 1, F to H).

We showed that multiple TFs and coregulators switch between the same two mobility states as H2B (Figs. 2 to 5). These data indicate that presumed ‘bound’ events can exhibit distinct mobility states. Using ligand-activated SRs, we determine that the lowest mobility state is associated with the active form of the TF (Fig. 2). PPARγ2 mutants show that chromatin engagement in state 1 requires an intact DNA-binding domain and an RXR heterodimerization domain (Fig. 5). To confirm that this state is associated with an active TF, we showed that over-expression of EGFP-C/EBPα, a TF that is known to cooperate with PPARγ2 at the chromatin level and facilitate its binding, leads to an increase in the proportion of PPARγ2 molecules in state 1.

Taken together, our data suggest a two-state model for chromatin and TF mobility. Chromatin is a viscoelastic polymer that has been shown to exhibit sub-diffusive dynamics (*17*). On our experimental timescales, chromatin explores a finite region of space we call a chromatin exploration domain (CED) (Fig. 7A). Within these CEDs, chromatin can exist in one of the two mobility states. On a timescale of 1.2 s, the lowest mobility state has an exploration diameter of ∼130-180 nm while the higher mobility state has an exploration diameter of ∼250-350 nm (Fig. 7, B and C). Chromatin can transition between these two mobility states due to processes yet to be determined. We have shown that TFs in their inactive form or with mutated DBDs and heterodimerization domains can primarily engage with chromatin in state 2. On the other hand, we find that active TFs can transition from state 2 chromatin to state 1 chromatin and vice versa.

**Fig. 7:**
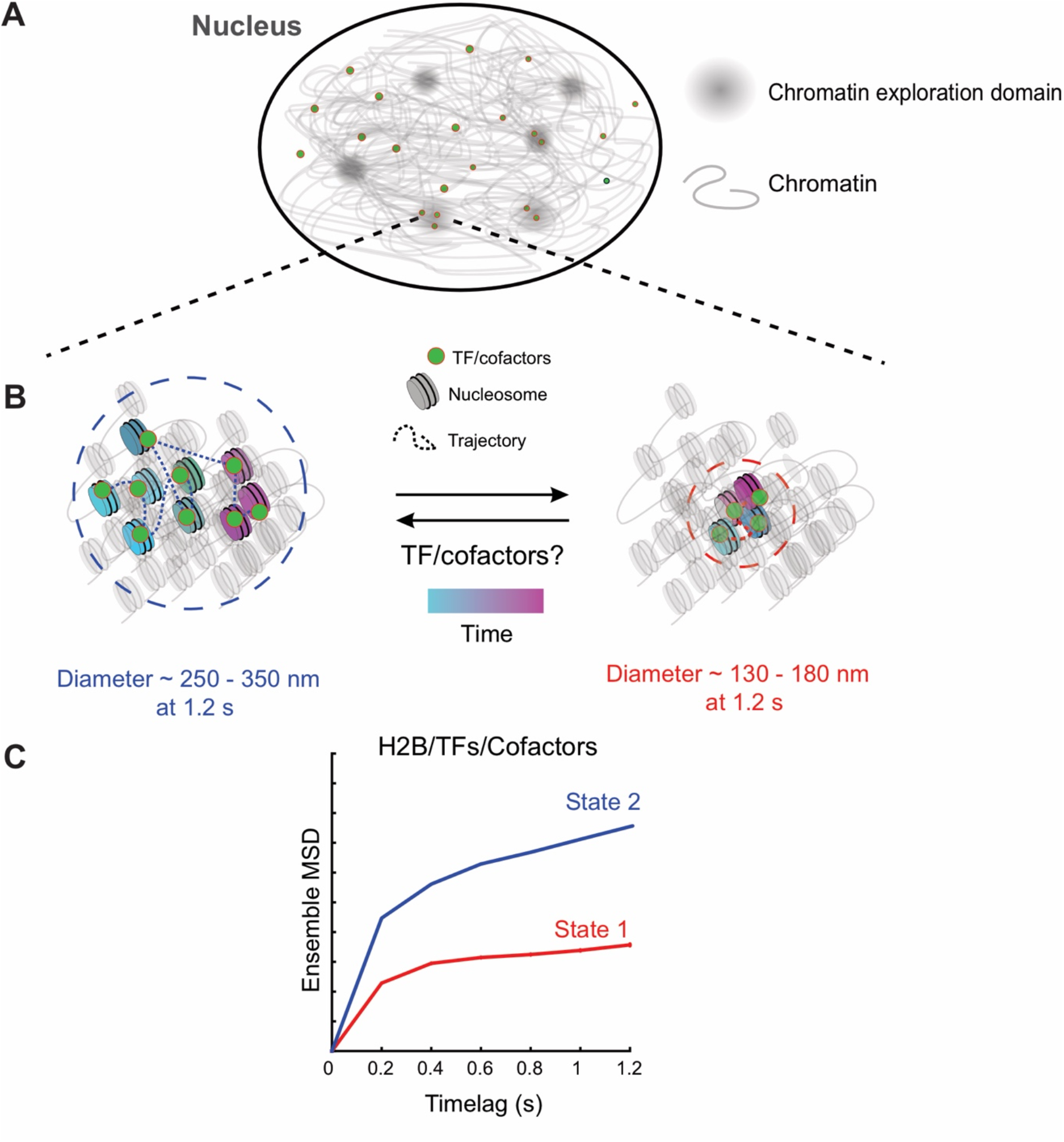
Two state model for chromatin and transcriptional regulators. **(A)** Over short timescales (∼ 1.2s), chromatin mobility is constrained within chromatin exploration domains. **(B)** Within these domains, transcription factors (TFs)/cofactors engage with chromatin and the TF-chromatin complex can exist in one of two mobility states. The left panel represents the higher mobility state (state 2), which has an exploration diameter of ∼250 nm – 350 nm at 1.2s. The trajectory shows the motion of a single TF/cofactor molecule over time. The lower mobility state (state 1) has an exploration diameter of ∼130 nm – 180 nm and the motion of a single TF/cofactor molecule is represented in the right panel. Chromatin and associated TFs/cofactors can dynamically switch between these two mobility states. TF binding can promote a switch from state 2 to state 1 or unbind from state 2 chromatin and bind to state 1 chromatin within the localized chromatin domain. **(C)** The mean squared displacement (MSD) plot of tracks classified by perturbation expectation maximization (pEMv2) is used to visualize the two different mobility states under the timescale of a single sub-track (1.2 s). The exploration diameter of the states is estimated as 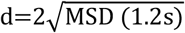.

The mobility state of TF that is interacting with chromatin can passively follow transitions in the mobility state of chromatin but whether TF binding can cause changes in chromatin mobility remains unclear. However, there is mounting experimental evidence in favor of this. RNA Pol II-mediated transcription has been shown to constrain nucleosome mobility (*7, 12*). Consistent with this result, TF binding and subsequent recruitment of the transcriptional machinery could trigger a transition of the local chromatin polymer (and of the bound TF) from state 2 to state 1 (Fig. 7B). Similarly, loop extrusion and nucleosome-nucleosome interactions have also been shown to constrain nucleosome mobility (*13*). However, directly establishing this will require advances in imaging to allow simultaneous tracking of a TF and a specific genomic locus at high spatial and temporal resolution.

We hypothesize that any transition to state 1 is the result of a combination of processes, comprising of but not limited to TF binding, RNA Pol II elongation, and loop extrusion. The following predictions emerge from this model. Inhibition of RNA Pol II with pharmacological drugs such as α-amanitin or DRB, both of which inhibit RNA Pol II elongation through different mechanisms, should result in an increase in the population fraction of state 2 and a reduction in the population fraction of state 1. Without our classification scheme, this would appear as an increase in the overall MSD of H2B, as recently reported (*12*). Similarly, rapid degradation of the RNA Pol-II subunit RPB1 or the cohesin complex subunit RAD21, using the auxin-inducible degron system would also result in an increase in the population fraction of state 2 relative to that of state 1 (*12, 13*).

As we have shown previously, TFs exhibit power-law distributed dwell times (*16, 20*). This broad distribution of dwell times renders it impossible to distinguish between specific and non-specific binding based on residence times alone. Different response elements are likely to present TFs with a broad affinity landscape. On the other hand, measuring the spatial mobility of TFs allowed us to identify two distinct mobility states across several classes of TFs in two cell lines. This opens new lines of inquiry hitherto unavailable through SMT.

Can we distinguish between specific and non-specific binding? The most elegant approach to answer this question would be to simultaneously track a single TF binding to a reporter gene along with transcriptional readout using an MS2- or PP7-stem loop system (*20*). Sparse labeling of TFs renders such events highly unlikely, but our data provide some clues to address this question. After classifying whole tracks into two states using the RL analysis, we find that tracks with overall lower mobility are preferentially in state 1 and exhibit much less switching than tracks with overall higher mobility (Fig. 6). If specific binding events occur preferentially within state 1, the long state 1 events in these low mobility tracks could represent specific binding while the transient state 1 events in the higher mobility tracks could represent TF hops from state 2 chromatin to state 1 chromatin.

In this study we have focused on long binding events. A long-standing question in the field is how TFs scan the 4D genome in search of their binding sites. TF motifs are typically 8-20 bp in length and are embedded within a sea of non-specific sequences (*5*). Theoretical considerations show that if TFs were to rely solely on Brownian motion to encounter their binding sites, they would take days to find a single specific binding site (*5*). Biophysical models of this apparent paradox suggest that bulk diffusion allows TFs to localize close to their specific sites, following which, they rely on 1D sliding, facilitated diffusion, and hopping to find their target motifs (*48, 49*). While these models are very provocative, little direct experimental evidence is currently available. As imaging technologies develop, and we push temporal and spatial resolutions to scales that are relevant for these processes, analysis tools presented here can help uncover modes of motion that remain elusive in conventional SMT studies. Applying these techniques to study TF dynamics in the context of development, disease, and evolution can provide a window into fundamental biological processes through the lens of individual transcription factors, paving the way for the development of targeted therapeutics for diseases driven by TFs gone awry.

## Limitations

To get a complete picture of TF dynamics from search to binding, we must be able to image with very high spatiotemporal resolution. Sparse labeling allows us to achieve sub-pixel localization, but our temporal resolution still suffers from photobleaching. We mitigate some of this by imaging with longer dark periods (200 ms) to capture long-lived binding events. However, this does not allow us to capture fast diffusing molecules since they diffuse out of the imaging volume on these timescales. MINFLUX tracking (*50, 51*) is currently the most promising nanoscopy technique that offers nanometer scale spatial resolution with a temporal resolution of hundreds of microseconds. Developments in fluorophore chemistry that improve the brightness and photo-stability of fluorophores will make longer imaging more feasible on instruments such as MINFLUX, and researchers will be able to interrogate both long- and short-time behaviors in the same set of tracks.

While our analysis provides evidence for two distinct mobility states in the nucleus, our MSD curves span only 6 timelags. With only 6 timelags, we cannot comment on the nature of the mobility states. To use the MSD to reliably distinguish between different physical models such as sub-diffusion, fractional Brownian motion, and confined diffusion, we need at least three decades of timelags (*52*). Non-MSD approaches to estimate diffusive parameters perform better than traditional MSD analyses but still require at least two decades of timelags (*53*). It is possible to achieve these long timescales by tracking sub-nuclear structures like telomeres, which can be labeled by the binding of multiple fluorescent proteins such as telomeric repeat factor 2 (TRF2) (*54*). However, photobleaching keeps these timescales outside the purview of SMT experiments. As can be seen from our analysis of long tracks, even with 200 ms dark periods, we can only span a 20-fold range of timelags.

Our study and all the SMT studies cited here (*12-15, 20, 21, 23, 24*) have been conducted in 2-dimensional cross-sections of the nucleus. It is possible for diffusing molecules to appear confined when projected in 2D. The higher mobility states recovered from pEMv2 for most TFs (colored green and yellow in all the figures) could represent a combination of this population of diffusive molecules along with tracking errors. This is supported by the fact that the proportion of these states is unchanged under all the perturbations. The only way to conclusively determine what these states represent will be to perform fast 3D tracking.

Finally, 2D tracking poses another significant challenge. When imaging molecules at the nuclear periphery or in perinucleolar regions, these molecules will undergo diffusion along an effective 2D surface. When these events are imaged in 2D, we are looking at the one-dimensional intersection of the surface and the focal plane. These events will preferentially appear to be in a very low mobility state since this is effectively one-dimensional motion. One must be careful to attribute these to the more compact nature of heterochromatin (*14*) without performing appropriate comparisons with 3D tracking.

## MATERIALS AND METHODS

### Cell lines and cell culture

3617 mouse adenocarcinoma cells (*25*) were grown in high glucose Dulbecco’s modified Eagle medium (DMEM, Gibco, #11960044) supplemented with 10% fetal bovine serum (FBS), 2 mM L-glutamine (Gibco #25030081), 1% MEM non-essential amino acids (Gibco #11140050), and 1 mM sodium pyruvate (Gibco, #11360070) at 37°C in a CO_2_ controlled incubator. 3617 cells contain stably integrated GFP-GR under a tetracycline-off system (*55*). To prevent the expression of GFP-GR, these cells were grown in the presence of 5 μg/mL of tetracycline.

3T3-L1 cells were cultured in DMEM supplemented with 10% calf serum (Gibco #26170043), 1% MEM non-essential amino acids, 1 mM sodium pyruvate, 50 U/mL penicillin and 50 μg/mL streptomycin (Gibco, #15070063) at 37°C in a CO_2_ controlled incubator.

### Plasmid constructs

#### H2B

pHalo-H2B was generated by PCR amplification of the H2B coding region from an H2B-GFP template and cloned into a pFC14A backbone (Promega, Madison, WI, USA) to fuse the HaloTag to the C-terminus of H2B (*56*).

#### Steroid receptors

The pHaloTag-GR plasmid expresses rat GR fused to HaloTag (Promega, Madison, WI, USA) in the C-terminus regulated by a CMVd1 promoter and has been described previously (*57*). pHalo-PR expresses human PR isoform beta fused with HaloTag at the N-terminus, regulated by a CMV promoter (*21*). pHalo-PR open reading frame (ORF) clone was purchased from Promega (Promega #FHC24423). pHalo-ER expresses human ERα fused to HaloTag in the C-terminus regulated by a CMVd1 promoter and has been described previously (*21, 58*). pHalo-AR expresses human AR with HaloTag fused to the C-terminus. This plasmid was custom-made by Promega and has been reported previously (*21*).

#### PPARγ2 and mutants

pHalo-PPARγ2 expresses human PPARγ isoform 2 fused to HaloTag in the N-terminus under a CMVd1 promoter (Promega ORF clone #FHC08305). PPARγ2 mutants were generated by nucleotide substitution using the QuikChange II XL Site Directed Mutagenesis Kit (Stratagene, La Jolla, CA, USA) following manufacturer’s protocol. PCR primers were designed using QuikChange Primer Design Program. All mutations were verified by sequencing.

#### Coregulators

pHalo-RELA expresses human NF-κB subunit p65 fused with HaloTag at the N-terminus in a pFN22K backbone. This construct was purchased from Promega. pHalo-GRIP1 expresses mouse GRIP1 with an N-terminus HaloTag fusion regulated by a CMVd1 promoter. This was generated by PCR amplification of the GRIP1 coding region from an EGFP-GRIP1 template and subsequent cloning into a pFN22K backbone using *SgfI* and *PmeI* restriction sites (*21*). pHalo-SMARCA4 expresses human SMARCA4 with HaloTag fused to the N-terminus under a CMVd1 promoter (Promega ORF FHC12075). pHalo-MED26 expresses human MED26 fused with a HaloTag at the N-terminus and was a kind gift from Joan Conaway’s lab. pHalo-CTCF expresses mouse CTCF with HaloTag fused to the C-terminus. This was generated by PCR amplification of the CTCF coding region from a CTCF-EGFP template (*59*) and cloned into the pHalo-GR backbone, which was cut using the *PvuI* and *XhoI* restriction enzymes (New England Biolabs, Ipswich, MA) and has been described previously (*16*).

#### EGFP construct

EGFP-C/EBPα expresses rat C/EBPα with an EGFP fusion on the N-terminus (this was a kind gift from Fred Schaufele, University of California San Francisco, San Francisco, CA, USA) and has been described previously (*60*).

#### Transient Transfections and agonist treatments

3617 and 3T3-L1 cells were plated in LabTek II (ThermoFisher, Waltham, MA, USA) or Cellvis (Mountain View, CA, USA) chamber slides for 24 hours before transfection. For 3617 cells, the indicated plasmids were transiently transfected using jetPRIME reagent (PolyPlus, New York, NY, USA) following manufacturer’s protocol. The protocol was optimized to prevent over-expression of HaloTag-protein chimeras (*21*). Cells were incubated in the jetPRIME reaction mixture containing 500 ng of DNA for 4 hours. The medium was then replaced with phenol red-free DMEM medium containing charcoal-stripped FBS (Life Technologies, Carlsbad, CA, USA) supplemented with 2 mM L-glutamine, 1% MEM non-essential amino acids, 1 mM sodium pyruvate, and 5 μg/mL tetracycline, and the cells were allowed to recover overnight.

For 3T3-L1 cells, 24 hours after plating, the medium was changed to optiMEM (Gibco, #31985070) and the cells were transfected with the indicated HaloTag- and/or EGFP-protein chimeras using Lipofectamine 2000 reagent (Invitrogen, Waltham, MA, USA) following manufacturer’s protocol. Briefly, for HaloTag-protein fusions, we used 750 ng DNA per 100 μL of Lipofectamine 2000 transfection mix. For EGFP-protein constructs, we used 4.5 μg of DNA per 100 μL of transfection mix. After incubating the cells in the transfection mix for 4 hours, the medium was replaced with fresh phenol red-free growth medium, and the cells were allowed to recover overnight.

Prior to imaging, the cells were incubated in medium containing 5 nM Janelia Fluor 549 (JF_549_) HaloTag ligand (*18, 61*) for 20 min. The cells were then washed three times with phenol red-free medium and returned to the incubator for 10 more minutes. Cells were then washed once more. 3617 cells were either left untreated or treated with 100 nM of the indicated hormone: dexamethasone (Dex), 17β-estradiol (E_2_), dihydrotestosterone (DHT), or progesterone (Prog) for 20 min before imaging. Dex, E_2_, DHT, and Prog were purchased from Sigma-Aldrich (St. Louis, MO, USA). 3617 cells expressing Halo-RELA were treated with 30 ng/mL of TNFα (Sigma-Aldrich, St. Louis, MO, USA) for 30 min before imaging. 3T3-L1 cells were all treated with 1 μM BRL49653/rosiglitazone (Rosi; Cayman Chemical Company, Ann Arbor, MI, USA) for 1 hour.

#### Microscopy

All samples were imaged on a custom-built HILO microscope in the LRBGE Optical Microscopy Core at NCI, NIH. Detailed information can be found in (*61*). Briefly, the microscope has a 150 × 1.45 numerical aperture objective (Olympus Scientific Solutions, Waltham, MA, USA); an Okolab stage-top incubator for temperature and 5% CO_2_ control (Okolab, Pozzuoli NA, Italy). The microscope is equipped with a 561 nm laser (iFLEX-Mustang, Excelitas Technologies Corp., Waltham, MA, USA) and an acousto-optical tunable filter (AOTFnC-400.650, AA Optoelectronic, Orsay, France) (*19, 61*). Images were collected using an EM-CCD camera (Evolve 512, Photometrics, Tucson, AZ, USA) every 200 ms (5 Hz frame rate) with an exposure time of 10 ms for a total of 2 min (600 frames) with a laser power of 0.96 mW (*16*).

#### Tracking

Particle detection and tracking are performed using TrackRecord v6, a custom tracking software written in MATLAB (version 2016a, The MathWorks Inc, Natick, MA, USA) that is publicly available at https://github.com/davidalejogarcia/PL_HagerLab/ and has been extensively described previously (*16, 20, 21, 56, 61*). The image stacks were filtered using top-hat, Wiener, and Gaussian filters. A hand-drawn region-of-interest (ROI) was used to demarcate the boundary of the nucleus. The particle detection intensity threshold was determined to be the lowest threshold at which less than 5% of detected molecules had a signal-to-noise ratio of 1.5 or less. Sub-pixel localization was achieved by fitting the detected particles to a two-dimensional Gaussian. Detected particles were then tracked using a nearest-neighbor algorithm (*62*) with a maximum allowed jump of 4 pixels, maximum allowed gap of 1 frame, and shortest track of 6 frames. Including motion blur, pEM estimates the localization precision to be ∼20 nm for state 1 and ∼40 nm for state 2 (*22, 63*). The higher mobility states have a localization precision of ∼70 nm.

#### Identification of distinct diffusive states using pEMv2

Perturbation-expectation maximization v2 (pEMv2) (*22*) was used to classify the single-molecule trajectories into multiple diffusive states. pEMv2 requires tracks to be divided into sub-tracks of equal length. We split our trajectories into sub-tracks of length 7 frames since longer tracks increase the likelihood of transitions within a sub-track. We ran pEMv2 independently on each protein and treatment to avoid forcing different datasets to converge on the same mobility states. No prior assumptions on the number of diffusive states or the types of diffusive motion were made (*22*). pEMv2 was allowed to explore between 1 and 15 states, with 20 reinitializations and 200 perturbations. The maximum number of iterations was set to 10,000 with a convergence criterion of 10^−7^ for the change in the log-likelihood function. Convergence of pEMv2 was verified through multiple runs. The covariance matrix was allowed to have three features.

After classification by pEMv2, each sub-track is assigned a posterior probability to belong to each of the states. For example, if pEMv2 converges to three states, each sub-track would have three posterior probabilities–one for each determined state. We assign each sub-track to the state for which it has the highest posterior probability (Fig. S1A).

A sub-track could have similar posterior probabilities to belong to two or more states. For instance, in our mock example with three states (Fig. S1A), we could have a sub-track with a posterior probability distribution of (0.9, 0.05, 0.05) in which case we would assign the sub-track to state 1. However, we could also have a sub-track with a posterior probability distribution of (0.5, 0.4, 0.1), in which case, while we would assign the sub-track to state 1, it has a very high probability to belong to state 2 as well. To mitigate this, we calculated ΔPP, which is the difference of the two highest posterior probabilities for each sub-track and excluded sub-tracks with ΔPP ≤ 0.2 from the ensemble MSD and population fraction calculations (Fig. S1B).

#### Calculation of the unbound fraction

States that account for less than 5% of all sub-tracks are excluded from the calculation of the population fraction (Fig. S1C). For consistent comparison of population fractions of steroid receptors before and after hormone treatment or PPARγ2 wildtype against mutants, we needed an estimate of the unbound fraction. Following the methodology outlined in (*16, 56*), we used the respective H2B jump histograms to calculate two jump distance thresholds for each cell line: R_min_ is the jump distance of 99% of H2B molecules between consecutive frames and R_max_ is the jump distance of 99% of H2B molecules between six frames (equal to the shortest track). Jump events larger than R_min_ over consecutive frames or larger than R_max_ over six frames were classified as unbound. For each species, the unbound fraction was then calculated as the ratio of the total number of unbound events to the total number of tracked molecules. For 3617 cells, R_min_ = 250 nm and R_max_ = 330 nm. For 3T3-L1 cells, R_min_ = 270 nm and R_max_ = 390 nm.

#### Transition probabilities

For the calculation of transition probabilities, since most of the tracks belong to low-mobility states 1 and 2, all the other states detected by pEMv2 were grouped together into a third “other” state. This allows us to calculate the transition probability among three states: low-mobility state 1, low-mobility state 2, and “other” states. For each track, the number of transitions between each pair of these states is calculated using a custom MATLAB script. Only tracks with at least three sub-tracks were included in this analysis. These transition counts are then added up to obtain a transition matrix *T* where the element *T*(*i, j*) is the number of transitions from state *i* to state *j*. This matrix is then normalized to obtain the transition matrix *P*_*t*_ where 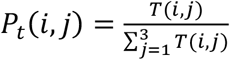.

To test whether these transition probabilities are different from those recovered from a randomized ensemble with the same population fraction, the sub-track state assignments are randomly shuffled, and the transition probabilities 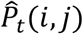 are calculated for this randomized ensemble. This process is repeated 1000 times and the statistical significance for a transition probability *P*_*t*_(*i, j*) is reported as the proportion of randomized trials with 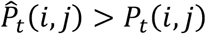.

### Estimation of the MSD distribution using the Richardson-Lucy algorithm

The single particle tracking data was used to calculate the self-part of the van Hove correlation function (vHc) as *G*_*S*_(*r, τ*) = *A*_*S*_⟨*δ*(*r*_*i*_ − |*r*_***i***_(*t* + *τ*) − *r*_*i*_(*t*)|⟩, where ***r***_***i***_ is the position of the i^th^ nucleosome and *A*_*S*_ = ∫ *d*^2^*rG*_*s*_(*r, τ*) is a normalization constant. The vHc is assumed to be a superposition of Gaussian functions, 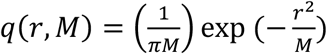 as *G*_*S*_(*r, τ*) = ∫ *P*(*M, τ*)*q*(*r, M*) *dM*, where *P*(*M*) is the distribution of mean-squared displacements of the population of nucleosomes. The Richardson-Lucy algorithm is used to extract *P*(*M*) from the empirical vHc as follows (*15*): from an initial distribution, 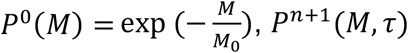 at the (n+1)^th^ iteration was iteratively obtained from

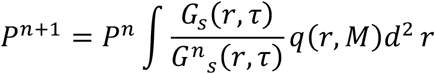

with the constraint that *P*^*n*^(*M, τ*) > 0 and normalized. The minima of *P*(*M, τ*) were used to classify individual nucleosome tracks into different mobility states.

## ACKNOWLEDGMENTS

We thank Tatiana S Karpova and David A Ball from the Optical Microscopy Core at the NCI, NIH for assistance in imaging and data processing, Luke Lavis (Janelia Farm Research Campus) for providing Halo-JF dyes, and Ivan Rey-Suarez (University of Maryland, College Park) for helpful discussions about the pEMv2 analysis.

## Author contributions

KW, DMP, AU, and GLH conceived the study. KW, DAS, RAMJ, VP, GF, and DMP performed the SMT experiments. RLS and RAMJ performed the PPARγ2 mutagenesis experiments and subsequent validation. KW developed the pEMv2 analysis workflow with input from AU. AU developed the RL analysis. KW wrote the original draft of the manuscript with input from DAS, DMP, AU, and GLH. All co-authors subsequently edited and reviewed the manuscript. DW and SM supervised the PPARγ2 section of the study including initial SMT analysis. DMP, AU, and GLH supervised the entire study.

## Data and materials availability

All data are available in the main text or the supplementary materials. Raw data, analysis scripts, and reagents are available at reasonable request from GLH

