## Supplementary Figures for "Single-molecule tracking reveals two low-mobility states for chromatin and transcriptional regulators within the nucleus"

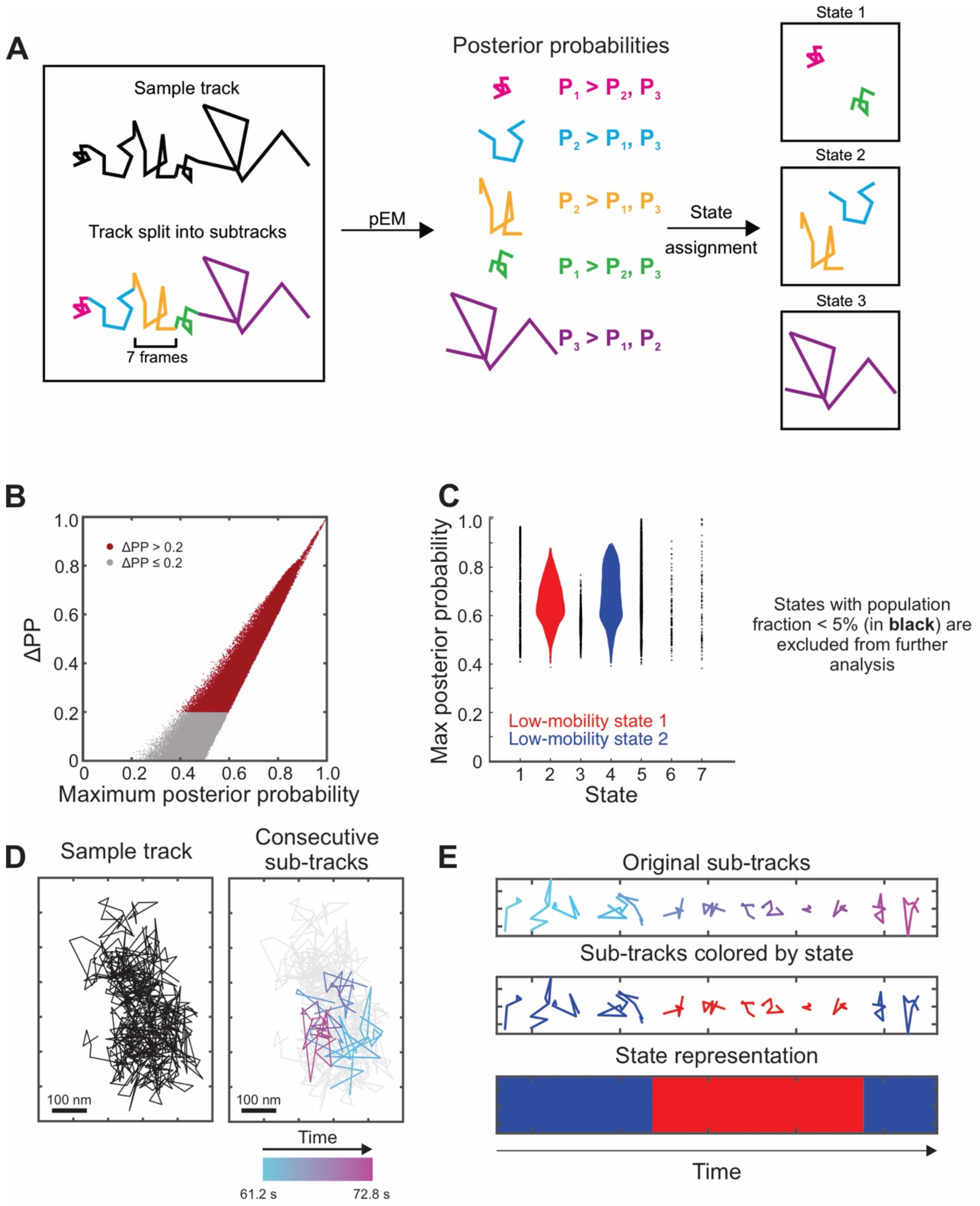


**Supplementary Figure 1**

**Fig. S1: pEM workflow and data representation. (A)** Cartoon showing pEM analysis workflow. Tracks are divided into sub-tracks of length 7 frames and classified into a state based on the posterior probability distribution. **(B)** Scatter plot of the difference of two highest posterior probabilities versus the maximum posterior probability. **(C)** Swarmchart of maximum posterior probability vs state (ordered in increasing order of diffusivity). States with population fraction less than 5% are colored in black and excluded from further analysis. **(D)** (left) Sample single molecule trajectory (right) Sub-tracks from 61.2s to 72.8s color-coded for time. **(E)** (top) Sub-tracks in panel D (right) separated in space for visualization (middle) Same tracks as in the top panel colored according to state assignment by pEMv2 (bottom) Block state representation to visualize state transitions. Each sub-track is represented by a red (state 1) or blue (state 2) pixel.

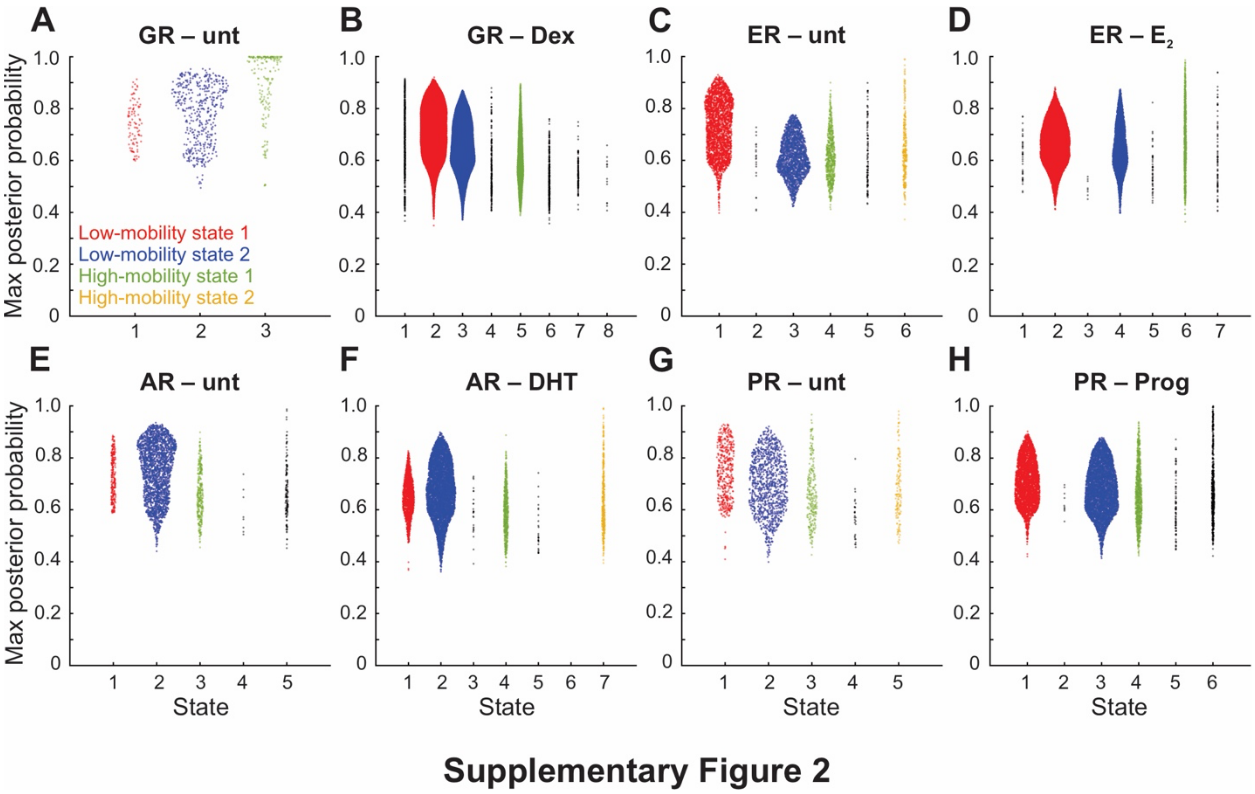


**Fig. S2: Swarmcharts of steroid receptors.** Maximum posterior probability vs pEMv2 state for **(A)** Glucocorticoid receptor (GR) without hormone, **(B)** GR activated with dexamethasone, **(C)** Estrogen receptor (ER) without hormone, **(D)** ER activated with 17β-estradiol ($E_2$), **(E)** Androgen receptor (AR) without hormone, **(F)** AR activated with dihydrotestosterone (DHT), **(G)** Progesterone receptor (PR) without hormone, **(H)** PR activated with progesterone (Prog).

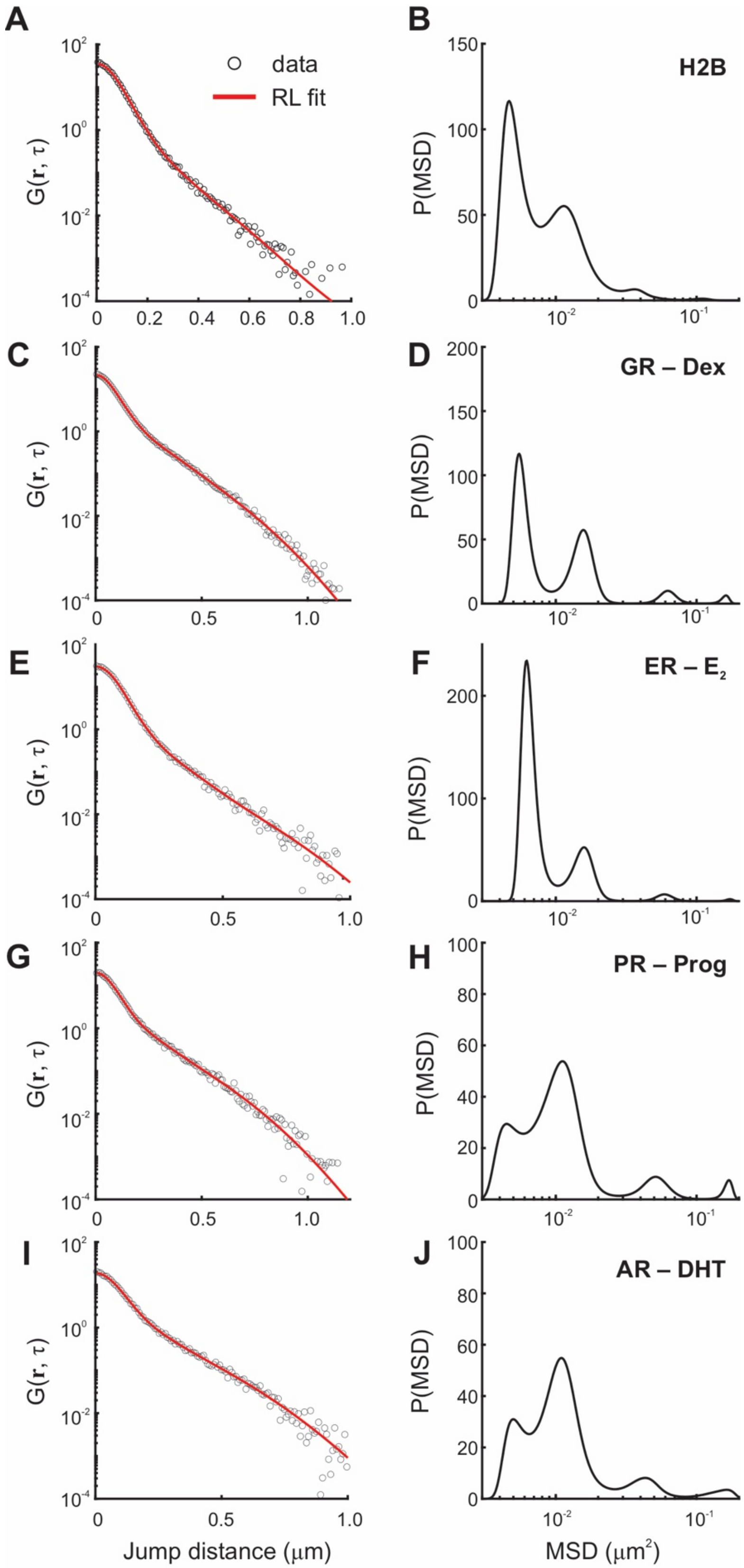


**Supplementary Figure 3**

**Fig. S3: Prediction of diffusivity distributions from van Hove correlations reveals distinct mobility states of transcription factors and histones**. **(A, C, E, G, I)** van Hove correlation function G(r,t) for single molecule trajectories of **(A)** H2B, **(C)** GR, **(E)** ER, **(G)** PR, and **(I)** AR calculated at t=0.8s. Red line is the computed vHc from P(MSD) using Richardson-Lucy inversion. **(B, D, F, H, J)** The distribution P(MSD) computed at 0.8 seconds for the molecular species indicated.

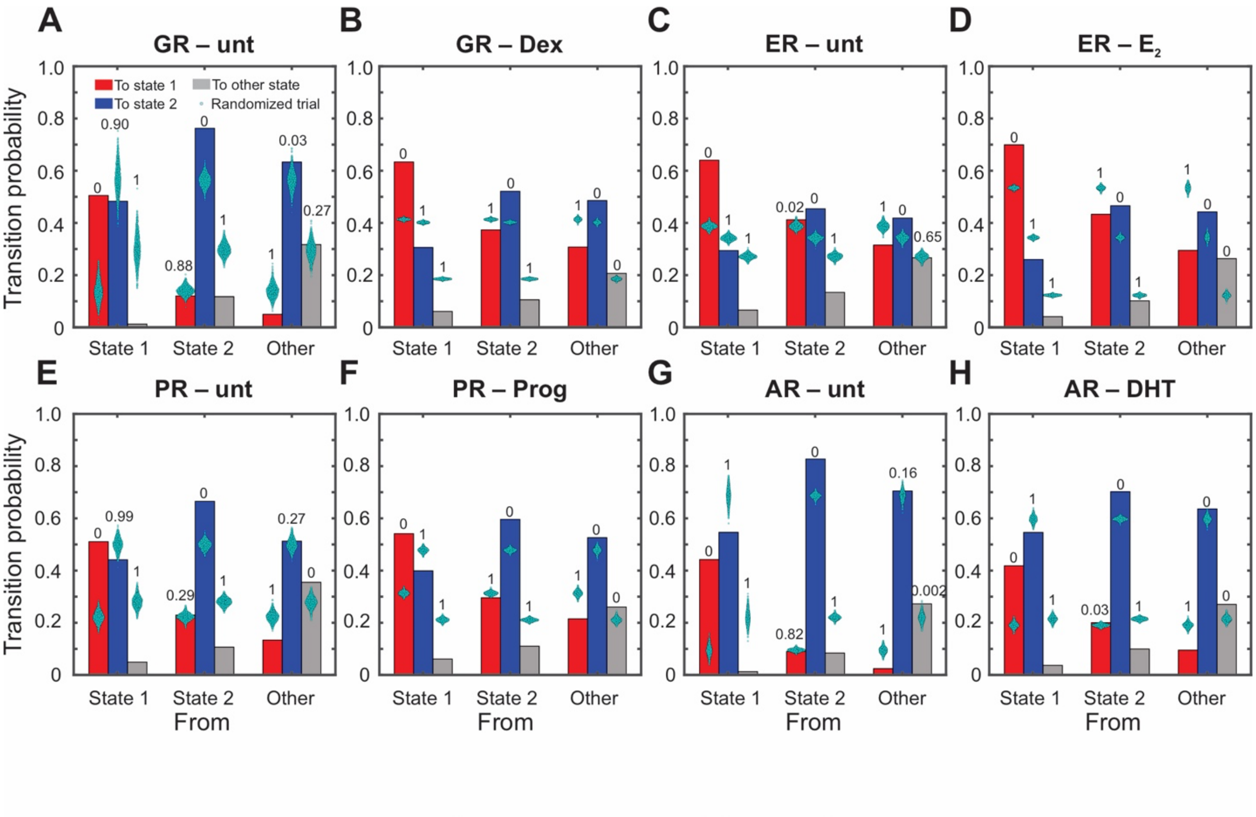


**Supplementary Figure 4**

**Fig. S4: Transition probabilities for steroid receptors compared against randomized trials.** Transition probabilities between states 1, 2, and "other". For simplicity, all the other states detected by pEMv2 were combined into a single state. The groups along the x-axis represent the transition probability from the indicated state while the colors of the bars represent transitions into state 1 (red), state 2 (blue), and other states (gray). Cyan swarmcharts represent transition probabilities calculated for 1000 randomized ensembles. Numbers above the bars indicate the proportion of randomized trials with transition probability higher than the corresponding calculated probability. **(A)** Glucocorticoid receptor (GR) – unliganded. **(B)** GR activated with dexamethasone (Dex). **(C)** Estrogen receptor (ER) – unliganded. **(D)** ER activated with 17β-estradiol ($E_2$). **(E)** Progesterone receptor (PR) – unliganded. **(F)** PR activated with progesterone (Prog). **(G)** Androgen receptor (AR) – unliganded. **(H)** AR activated with dihydrotestosterone (DHT).

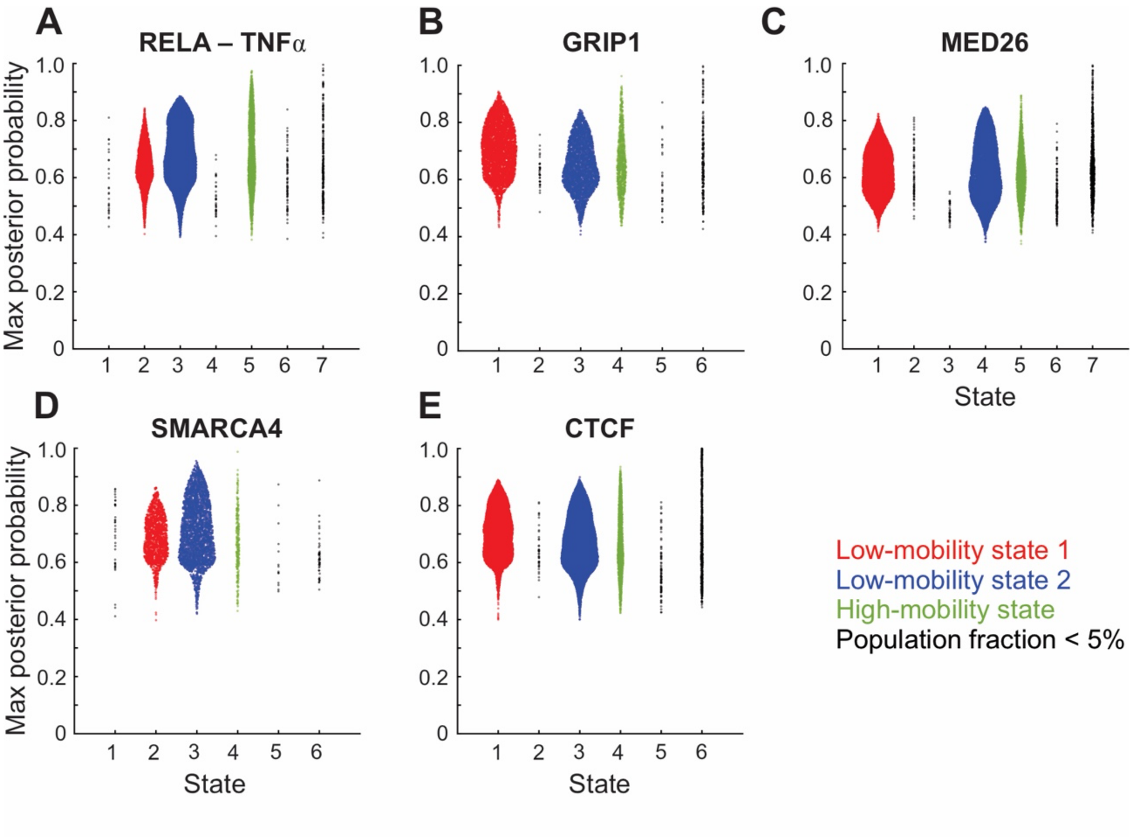


**Supplementary Figure 5**

**Fig. S5: Swarmcharts of maximum posterior probability as a function of states identified by pEMv2.** Maximum posterior probability vs pEMv2 state for **(A)** RELA activated with TNFα, **(B)** GRIP1, **(C)** MED26, **(D)** SMARCA4, **(E)** CTCF.

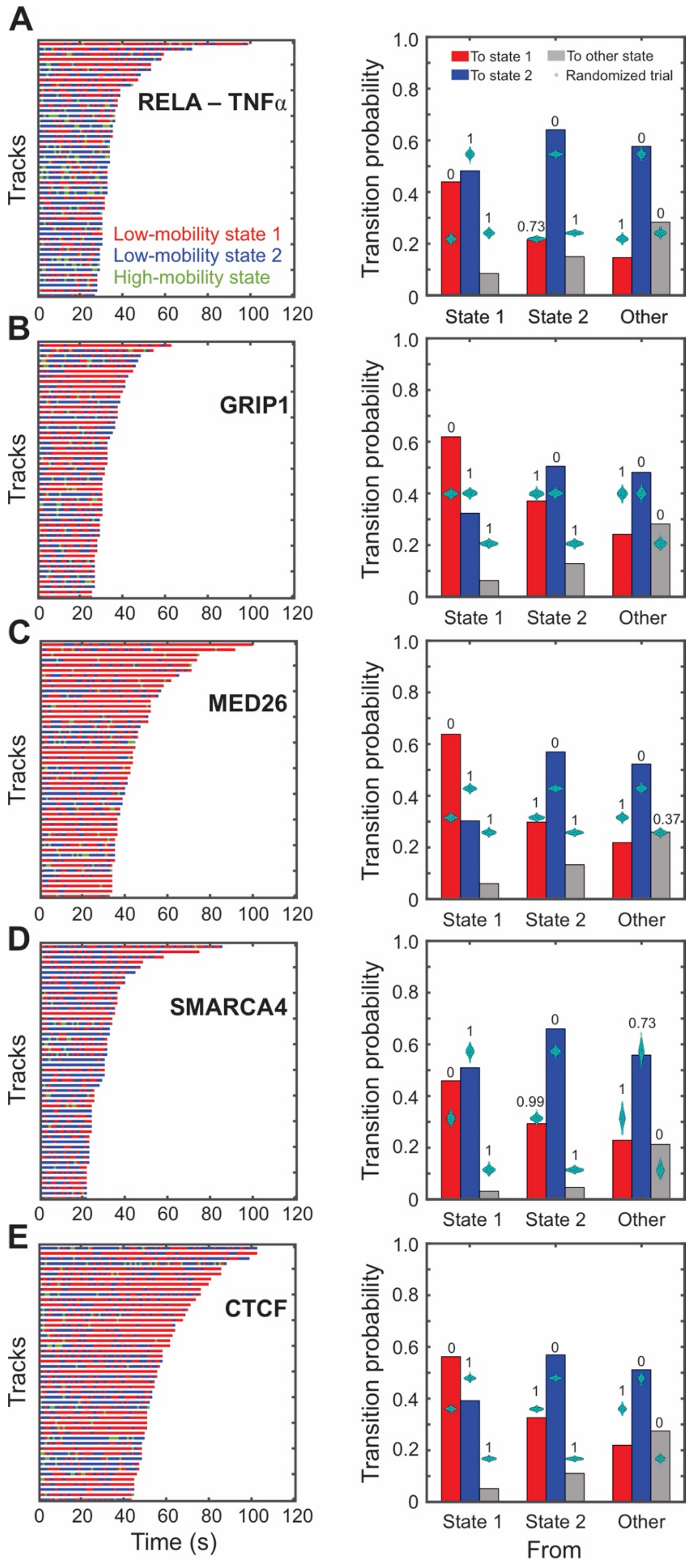


**Supplementary Figure 6**

**Fig. S6: All tested coregulators exhibit switching between mobility states.** For all panels: (left) Temporal reconstruction for 50 longest tracks color-coded by state assignment. (right) Transition probabilities between states 1, 2, and “other”. For simplicity, all the other states detected by pEMv2 were combined into a single state. The groups along the x-axis represent the transition probability from the indicated state while the colors of the bars represent transitions into state 1 (red), state 2 (blue), and other states (gray). Cyan swarmcharts represent transition probabilities calculated for 1000 randomized ensembles. Numbers above the bars indicate the proportion of randomized trials with transition probability higher than the corresponding calculated probability. **(A)** RELA treated with TNF$\alpha$, **(B)** GRIP1, **(C)** MED26, **(D)** SMARCA4, **(E)** CTCF.

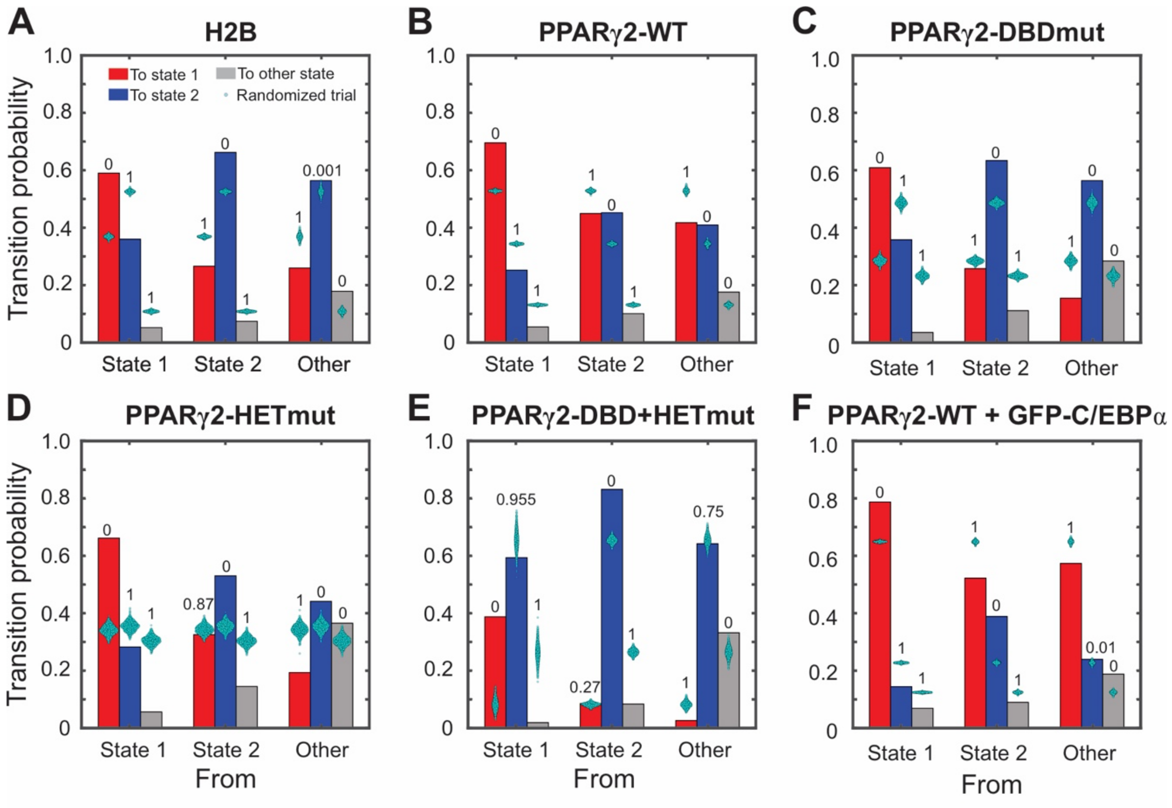


**Supplementary Figure 7**

**Fig. S7: Transition probabilities for PPARγ2 mutants and H2B in 3T3-L1 pre-adipocytes**. Transition probabilities between states 1, 2, and "other". For simplicity, all the other states detected by pEMv2 were combined into a single state. The groups along the x-axis represent the transition probability from the indicated state while the colors of the bars represent transitions into state 1 (red), state 2 (blue), and other states (gray). Cyan swarmcharts represent transition probabilities calculated for 1000 randomized ensembles. Numbers above the bars indicate the proportion of randomized trials with transition probability higher than the corresponding calculated probability. **(A)** H2B, **(B)** PPARγ2-WT, (C) PPARγ2-DBDmut, **(D)** PPARγ2-HETmut, **(E)** PPARγ2-DBD+HETmut, **(F)** PPARγ2-WT + GFP-C/EBPα.

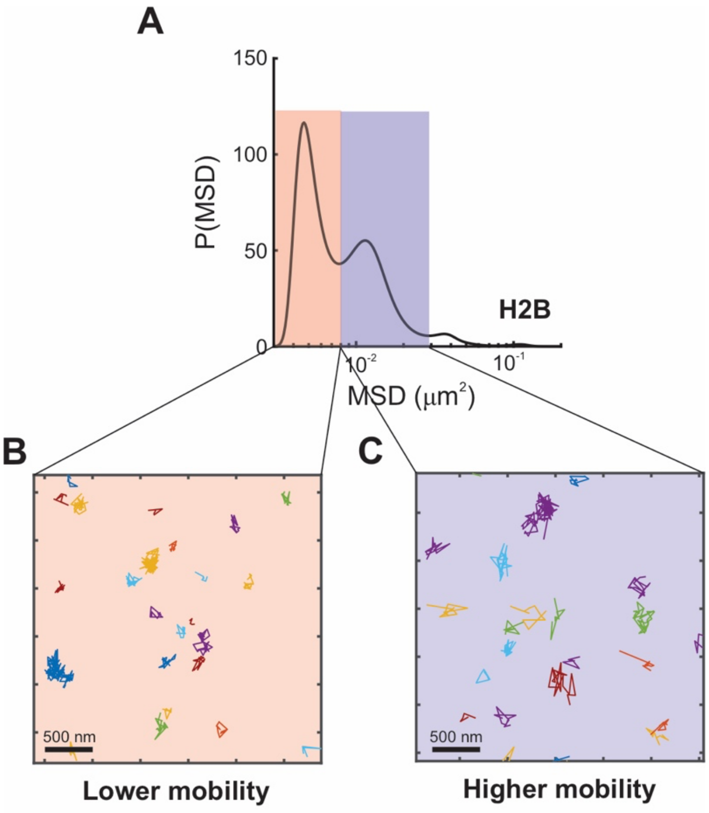


### Supplementary Figure 8

**Fig. S8: Classifying tracks based on MSD distribution. (A)** Minima in the predicted mean-squared distribution (MSD) distribution can be used to classify trajectories into two groups: tracks with an overall lower and higher mobility at 0.8 s. H2B tracks with an MSD in the red region are classified into the lower mobility group while those with an MSD in the blue region are classified into the higher mobility group. **(B)** Representative trajectories classified into the lower mobility group. **(C)** Representative trajectories classified into the higher mobility group.

**Movie S1.**
Representative single-molecule tracking movie of an H2B-Halo expressing 3617 cell. Images are collected every 200 ms with 10 ms exposure. Scale bar 4 μm.
